# Systematic mapping of antibiotic cross-resistance and collateral sensitivity with chemical genetics

**DOI:** 10.1101/2024.01.25.576750

**Authors:** Nazgul Sakenova, Elisabetta Cacace, Askarbek Orakov, Florian Huber, Vallo Varik, George Kritikos, Jan Michiels, Peer Bork, Pascale Cossart, Camille Goemans, Athanasios Typas

## Abstract

By acquiring or evolving resistance to one antibiotic, bacteria can become resistant to a second one, due to shared underlying mechanisms. This is called cross-resistance (XR) and further limits therapeutic choices. The opposite scenario, in which initial resistance leads to sensitivity to a second antibiotic, is termed collateral sensitivity (CS) and can inform cycling or combinatorial treatments. Despite their clinical relevance, our current knowledge of such interactions is limited, mostly due to experimental constraints in their assessment and lack of understanding of the underlying mechanisms. To fill this gap, we used published chemical genetic data on the impact of all *Escherichia coli* non-essential genes on resistance/sensitivity to 40 antibiotics, and devised a metric that robustly discriminates between known XR and CS antibiotic interactions. This metric, based on chemical genetic profile (dis)similarity between two drugs, allowed us to infer 404 XR and 267 CS interactions, thereby expanding the number of known interactions by more than 3-fold – including reclassifying 116 previously reported interactions. We benchmarked our results by validating 55 out of 59 inferred interactions via experimental evolution. By identifying mutants driving XR and CS interactions in chemical genetics, we recapitulated known and uncovered previously unknown mechanisms, and demonstrated that a given drug pair can exhibit both interactions depending on the resistance mechanism. Finally, we applied CS drug pairs in combination to reduce antibiotic resistance development in vitro. Altogether, our approach provides a systematic framework to map XR/CS interactions and their mechanisms, paving the way for the development of rationally-designed antibiotic combination treatments.

## Introduction

While the spread of antibiotic resistance is increasing at alarming rates^1^, fewer and fewer novel antibiotics are being approved for clinical use^2,3^. Importantly, the development of intrinsic or horizontally acquired resistance to a given drug can lead to cross-resistance (XR)^4^ to other drugs, limiting treatment options. The same processes can also give rise to collateral sensitivity (CS)^5^ to other drugs, due to trade-offs or fitness costs of resistance mechanisms^6,7^ (**Fig. 1a**). The principle of CS has been successfully used to reduce the rates of resistance emergence^8–15^, or to even re-sensitize microbes to antibiotics^16^, by combining or cycling of CS drug pairs. The benefits of avoiding the use of XR drug pairs in tandem or consecutively are obvious. Overall, it is imperative to systematically map and understand XR and CS relationships between drugs, especially in an era of diminishing therapeutic options.

**Fig 1.**
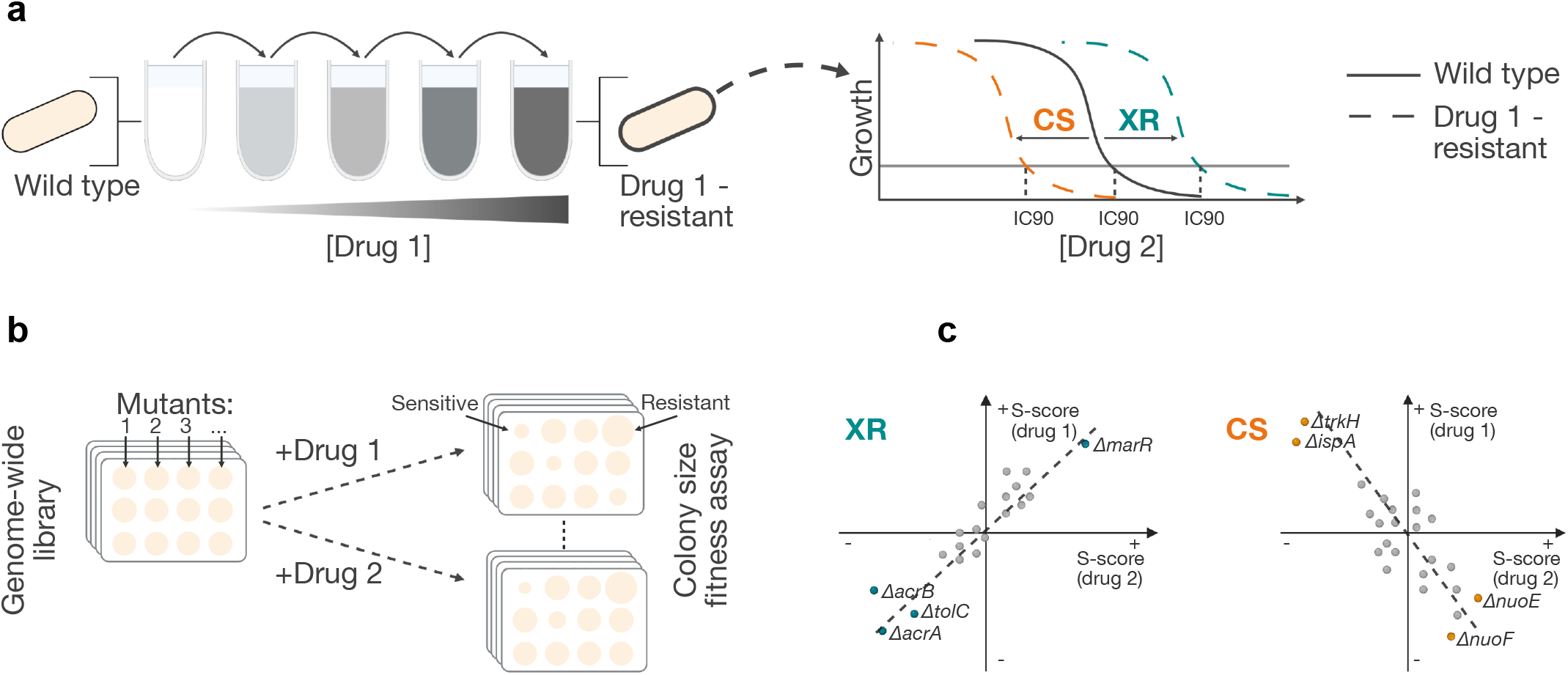
Chemical genetics allows for systematic XR and CS assessment. **a**, Schematic illustration of conventional way to assess XR/CS drug interactions via experimental evolution. Resistant mutants are raised against drug 1 and then tested for susceptibility to drug 2. The MIC/IC_90_ is compared to that of the ancestral strain. **b,** Schematic illustration of chemical genetic screens with arrayed libraries. Several drugs (1, 2 …) are profiled across genome-wide gain-of-function or loss-of-function mutant libraries. The fitness of each mutant is evaluated independently, e.g. by measuring colony size. **c,** XR and CS are associated with chemical genetic profile similarity and dissimilarity, respectively. The fitness of deletion mutants (s-scores; positive and negative scores denote increased and decreased fitness respectively) is plotted for two drugs simulating XR and CS paradigms in *E. coli*. Labelled mutants are involved in known mechanisms of XR and CS^17–19^. If the same mutations make cells more resistant or sensitive to two drugs, cells are more likely to evolve mechanisms that inhibit or promote these exact processes during evolution and become XR to both drugs. The opposite is true for CS.

The most common approach to measure XR and CS is to experimentally evolve resistance against one drug, and then to measure susceptibility to another drug for a number of evolved lineages (**Fig. 1a**). Our understanding of the underlying mechanism(s) relies on sequencing the genomes of the evolved strains to identify recurrent genetic alterations^8,17–19^. Although powerful, this approach has limitations in terms of effort, scale and costs. Hence current knowledge of XR and CS interactions is limited to a few bacterial species and a small number of antibiotics^8,9,12,16–26^. Importantly, experimental evolution probes a limited number of lineages and a small part of the solution space in terms of possible resistance mutations, which strongly depends on selection pressure applied. This may lead to inconsistencies when assessing drug pair interactions. Furthermore, evolution experiments inevitably lead to numerous additional mutations that make the mapping of causal resistance mechanisms difficult without additional experiments. To facilitate drug susceptibility testing of experimentally evolved strains or to dissect the evolved resistance mechanism(s), adaptations to the original method have been proposed, e.g. the automation of minimum inhibitory concentration (MIC) measurements^26^, or the phenotypic characterization of evolved strains with transcriptomics^27,28^. Even though these adaptations increase the number of lineages, chemicals and interactions probed, they still explore a limited genetic space for resistance, and require extensive sequencing and prior knowledge to identify the causal resistance mechanisms. Here, we set out to overcome these limitations by developing a predictive, sequencing-free framework based on drug-gene interactions, and harnessing the systematic nature of chemical genetic screens.

Chemical genetics involve the systematic assessment of drug effects on genome-wide mutant libraries^29,30^. Such data have been previously shown to capture information on drug mode of action, resistance and interactions in *E. coli*^31–36^. Importantly, chemical genetics systematically quantify how each gene in the genome contributes to resistance or susceptibility to a large set of drugs. The first large-scale chemical genetics study assessed close to 50 antibiotics over different concentrations in *E. coli*, along with several other conditions (including other antimicrobial compounds and non-antibiotic drugs) against an arrayed library of 3979 single-gene deletion mutants or alleles^31^ (**Fig. 1b**). The similarity between chemical genetic profiles for different drugs has been reported to correlate with XR frequency^18^, and has been used to minimize XR between antimicrobial peptides and antibiotics^37^. Several years ago, we proposed that such chemical genetic data would have in principle the capacity to identify both XR and CS interactions by comparing drug profiles^30^ (**Fig. 1c**), expediting the systematic identification of XR/CS interactions and mapping of their underlying mechanisms.

In this study, we used available *E. coli* chemical genetics data^31^ for 40 antibiotics (**Methods**) and explored different similarity metrics to identify the one best discerning between known XR and CS interactions. We applied this metric to many more drug pairs than probed collectively before, discovering three times more XR and six times more CS interactions than previously identified, including the reclassification of 116 previously wrongly reported drug-pair relations. We independently validated 7% (59/840) of these interactions by experimental evolution with 93% accuracy. By integrating all data into a drug-interaction network, we could examine the monochromaticity (i.e. if a given interaction is exclusively XR or CS) and conservation within antibiotic classes, identifying antibiotic (classes) with extensive XR or CS interactions. Next, we took advantage of the available chemical genetics data to track back the mutations responsible for specific interactions, thereby confirming known and resolving new mechanisms experimentally. Lastly, we showed that newly identified CS pairs used in combination could reduce resistance evolution compared to single drugs. Overall, we present a framework to accelerate XR/CS discovery and mechanism deconvolution, paving the way for rationally designed combinatorial, cycling, or sequential antibiotic treatments.

## Results

### Building a training set of known XR/CS interactions from evolution experiments

To build a training set of known XR/CS interactions, we collected data from four studies that performed experimental evolution in *E. coli*^8,17–19^. The majority of interactions (78%-338/429) had only been tested in one study. From the 91 antibiotic pairs tested in at least two studies, only a third (n=30; 20 Neutral, 9 XR and 1 CS) received the same assessment across studies, whereas 56 were called XR or CS interactions in one study, but neutral in the other (**Fig. 2a**). This suggests that XR/CS detection *via* experimental evolution is prone to high error rates, which could be due to several reasons: selection biases in evolution experiments (e.g. different selection pressure, drug resistance level cutoffs), slightly different criteria used in each study to define XR/CS, low power to call interactions (limited number of lineages tested), and population complexity (resistance or sensitivity assessment is done for lineage populations). We reasoned that most errors were likely due to false negatives, as studies were under-sampling the antibiotic resistance solution space. For this reason, we designated as XR or CS drug pairs that exhibited an interaction in at least one study, even if they were neutral in other(s). In contrast, drug pairs displaying conflicting responses, that is XR in one study and CS in another, were excluded (n=5). After comparing to drugs for which chemical genetic data is available^31^, we came up with 206 drug pairs (111 neutral, 70 XR and 25 CS), involving 24 different antibiotics (**Source Table of Fig. 2**). In chemical genetics, the drug effects on each mutant are represented as s-scores – those assess the fitness of a mutant in one condition, normalized by its fitness across all conditions^38,31^ (Methods, **Supplementary Table 1)**.

**Fig 2.**
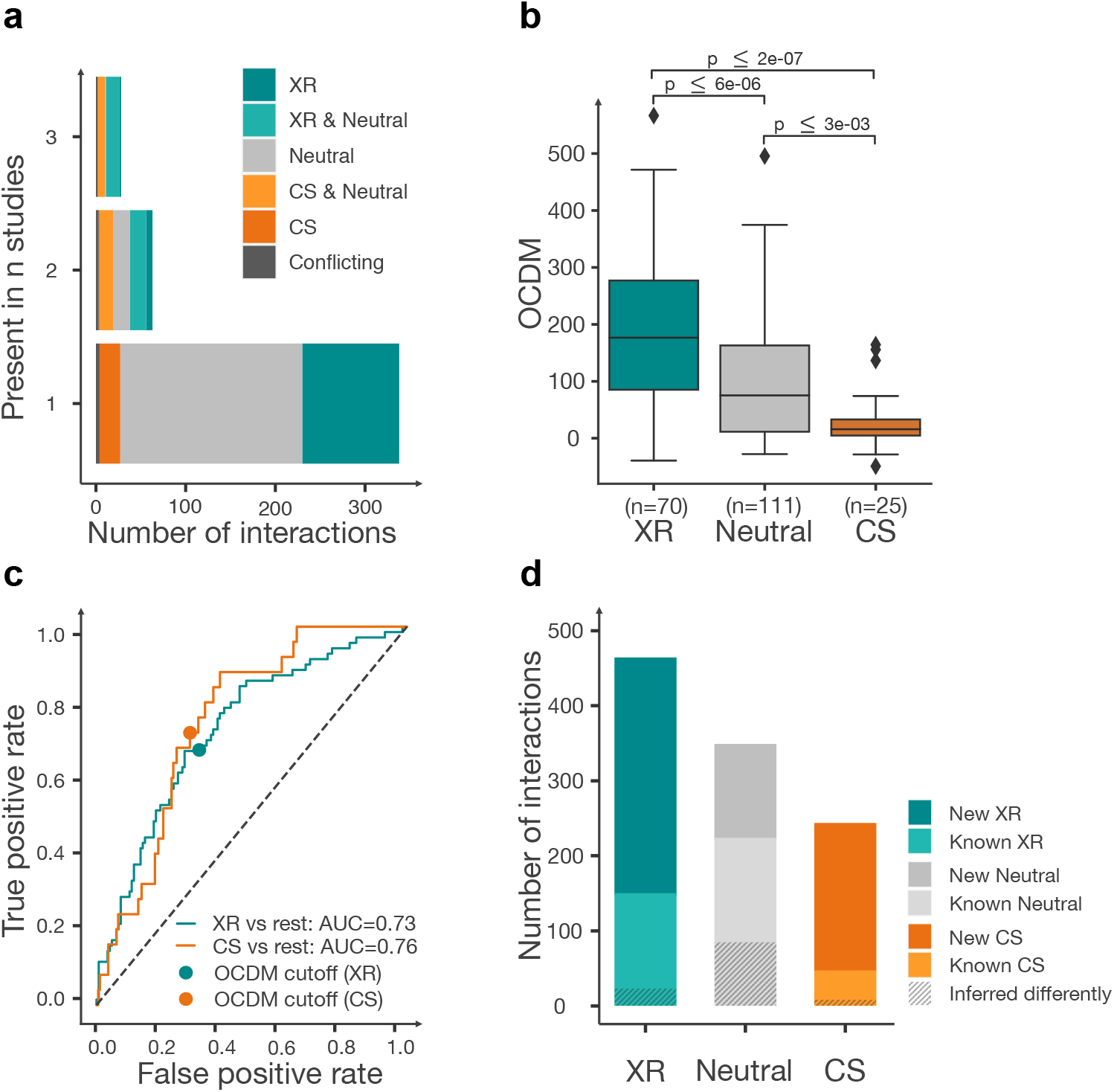
Chemical genetics-derived metric separates well known XR and CS interactions, and infers new ones. **a**, The overlap between published XR/CS interactions from four existing datasets^8,17–19^. is low. **b,** A devised metric derived from chemical genetic profile similarity, OCDM, can robustly discern between known XR, CS, and neutral interactions. P-values were obtained from two-sided paired Mann-Whitney U test. **c,** Receiver operating characteristic (ROC) curves for classification of XR (positive class) versus non-XR (negative class) and CS (positive class) versus non-CS (negative class). Each OCDM cutoff represents a point on the curve and is associated with a true positive rate and a false positive rate. The OCDM cutoffs chosen for XR and CS interactions are depicted with a circle. **d,** New XR, neutral and CS pairs inferred by chemical genetics using the OCDM cutoff are 2- and 4-fold more that currently known XR and CS interactions in *E. coli*. This difference is actually larger, as we reclassify 27.6% (n=116) of the known interactions (**Extended Data Fig. 2**). Known interactions include those between drugs for which there is no available chemical genetics (total n=420).

### Chemical genetic profile concordance identifies XR and CS drug pairs

Using our training set, we hypothesized that drugs sharing resistance mechanisms (XR) should have concordant chemical genetic profiles (i.e. most *E. coli* mutants would behave similarly when treated with each drug), as previously suggested for a subset of XR pairs (n=36)^18^. The opposite should be true for CS pairs, as mutations that would cause resistance to one drug would sensitize cells to another drug, leading to discordant chemical genetic profiles for the two drugs (**Fig. 1c**). We used different metrics derived from chemical genetic data to test whether we could discriminate between known XR, CS or neutrality (Methods). First, we assessed metrics of correlation between chemical genetic profiles, which exhibited low performance in discriminating between known XR, CS and neutral interactions (Area Under the Receiver Operating Curve: 0.52-0.67; **Extended Data Fig. 1a**). We reasoned that the noise generated by the high proportion of neutral phenotypes in chemical genetic data^31^ was compromising performance. To overcome this, we used six features based only on extreme s-scores per condition: sum and count of positive concordant s-scores, of negative concordant s-scores, and of total discordant s-scores (**Methods**). We then trained several machine learning classifier models (decision tree models) with these features for each drug pair. Such a trained classifier performs well, with F1 score, recall, precision and AUC ROC consistently exceeding 0.7 (**Extended Data Fig. 1b**). To avoid overfitting of a model based on a training dataset of XR/CS with caveats described before (**Fig. 2a**), we aimed to interpret the model instead of applying it directly on our test dataset. We learned from decision tree attributes (**Extended Data Fig. 1c**) that the sum and count of concordant negative s-scores are the most informative features, followed by the sum of discordant s-scores. Additionally, if the count of concordant negative s-scores is higher than the median count of concordant hits (this is mutants showing extreme positive or negative s-scores in both drugs) across all drug pairs (which is 7), the level of discordance is not important to classify interactions. Placing these attributes in an experimental evolution setting, this means that presence of mutants with resistance to both drugs (concordance) in heterogeneous populations would result in XR, while presence of only discordant mutants would lead to CS. Using this information we generated an Outlier Concordance-Discordance Metric (OCDM) that can discriminate between previously reported CS and XR interactions from the rest (AUC = 0.76 and 0.73, respectively; **Fig. 2b-c**; **Source Table of Fig. 2**, **Methods**), and selected the cutoff for extreme s-scores based on OCDM performance (**Extended Data Fig. 1d**). We then used the OCDM cutoffs **(Fig. 2c; Methods**) to classify all possible interactions between the 40 antibiotics within the chemical genetics data^31^. This yielded 634 new drug pair relationships (313 XR, 196 CS, 125 neutral), expanding the number of currently known XR and CS interactions by two and four times, respectively (**Fig. 2d**; **Supplementary Table 2**). In terms of previously measured drug pairs (n=206), our metric agreed for 90 and disagreed for 116 with previous calls, the latter coming mostly from previously called neutral interactions (**Extended Data Fig. 2a-c**), and thus potential false negatives. This increased the total number of inferred drug pair relationships to 840 (404 XR, 267 CS, 169 neutral), and expanded the number of known XR and CS interactions by three and six times, respectively (**Extended Data Fig. 2d**).

### Chemical-genetics based metric detects XR and CS with high accuracy

To benchmark our chemical genetic based metric (OCDM) and cutoff decisions, we selected a subset of 35 newly inferred and 24 previously tested drug pairs (for 13/24 we predicted a different interaction than one previously reported), and measured their interactions with experimental evolution. In our experimental evolution setup, we evolved resistance to 23 antibiotics in 12 parallel lineages, and tested resistant lineages for changes in susceptibility to a second antibiotic (**Supplementary Table 3**; **Fig. 3a**; **Methods**). Drug pairs were chosen to cover a wide OCDM range and to have low initial MICs to be able to evolve several-fold resistance. Pairs of antibiotics belonging to the same chemical class were mostly excluded from the validation set to avoid inflating the prediction accuracy of XR predictions, as such drug pairs are highly likely to be XR because of common resistance mechanisms. Evolving resistance to both drugs of each pair allowed us to assess whether interactions are (bi)directional, something we did not account for in the OCDM score. XR interactions are by definition bidirectional (at least to a certain) degree, and failure to detect them both ways in experimental evolution experiments exemplifies the limitation of the method. In contrast, CS interactions can be directional, as resistance mechanisms for each drug of the pair are different, and do not have to bear a fitness cost in the other drug. As a consequence, most of the previously detected CS pairs have been unidirectional. Furthermore, evolving resistance to 12 lineages allowed us to gain insight into the monochromaticity of interactions, that is whether drug pairs showed exclusively one type of interaction.

**Fig 3.**
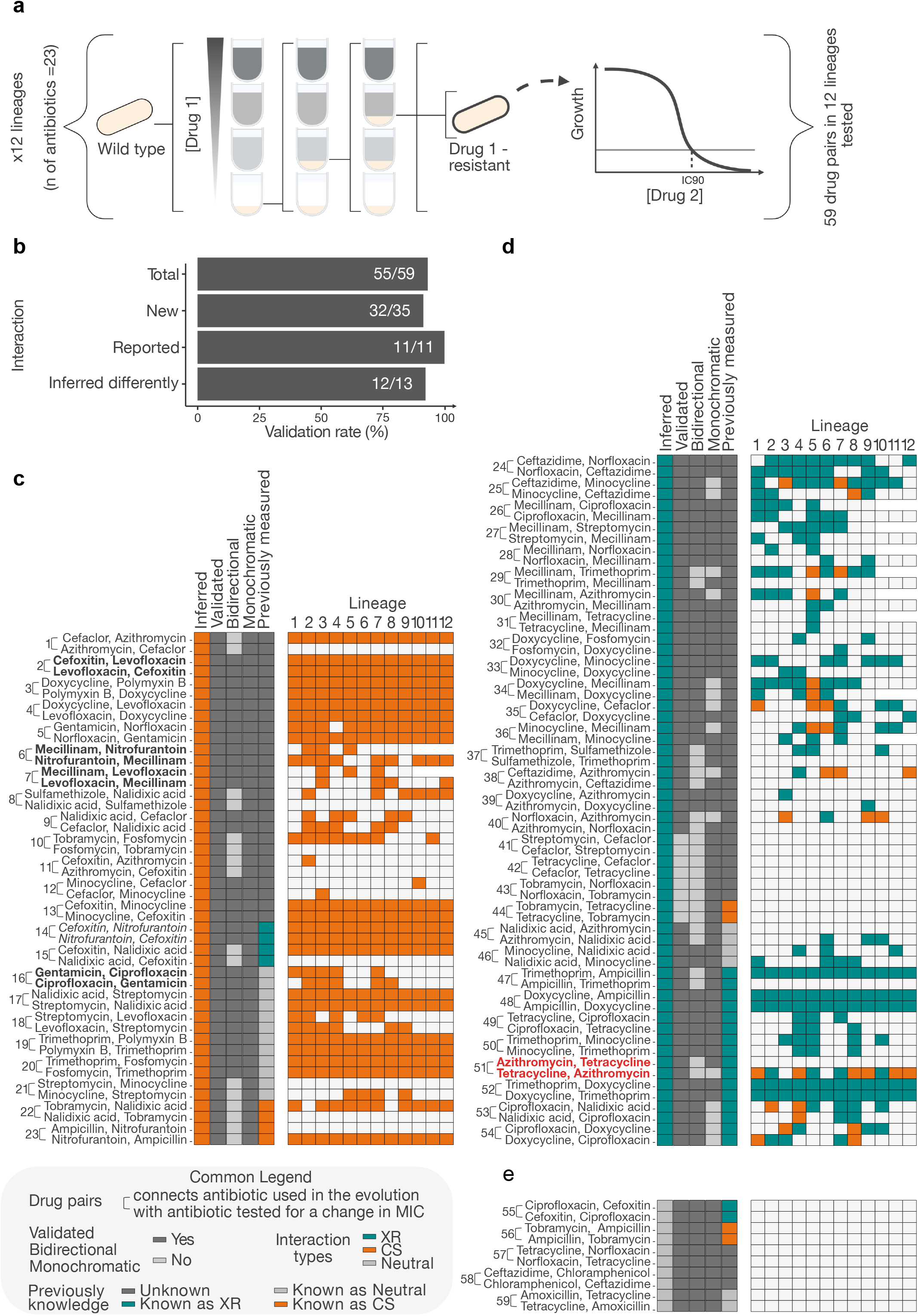
Inferred XR/CS interactions are validated with high accuracy. **a**, Schematic of benchmarking done by experimental evolution and MIC measurement for 59 drug pairs. Twelve lineages are evolved in parallel for 5 passages in 23 antibiotics. In each passage the culture growing at the highest concentration is transferred. The MIC of the final resistant population is then measured for all lineages in the relevant antibiotics. **b,** Most inferred interactions are experimentally validated, whether those are previously known (and our inference agreed/disagreed; latter designated as reclassified) or new. We considered an interaction to be validated if at least one lineage had log_2_ MIC fold-change > 1 for XR and < - 1 for CS, compared to the wildtype. **c-e,** Heatmap of 59 new, known (positive control), and reclassified interactions, split depending on whether they were inferred as CS (**c**), XR (**d**) or neutral (**e**). Interactions were tested in both directions, and directions are shown one after the other - the drug for which selection occurred is shown first, and the drug for which MIC was tested comes second. In each interaction, all tested lineages are shown. Interaction monochromaticity (whether interaction is exclusively CS or XR - neutral lineages do not affect this call), and published interaction assessment are also shown. Reclassified interactions are those for which our inference and validation agree, but previous reports have missed or reported wrongly. Interaction in red (least monochromatic interaction) is used in **Fig. 5** to understand the mechanisms in play. Interactions in bold are used later in **Fig. 6** to test resistance evolution in drug combinations. The interaction in italics (drug pair #14), which was conflicting across studies (XR in one study and CS in another), has been inferred and validated to be CS.

In total, we validated all but four of the inferred interactions, amounting to a total validation rate of 93.2%, and 91.4% of newly inferred interactions (**Fig. 3b**; **Source Table of Fig. 3**). Not only did we confirm those interactions for which literature and our metric agreed (n=11), but also 12 out of 13 interactions for which our predictions contradicted previous studies (**Fig. 3b**; **Extended Data Fig. 2**). These included 8 false negative (6 CS and 2 XR, reported neutral before) and 4 false positive (as 1 XR and 3CS) cases (**Fig. 3c-e**). This highlights the superior accuracy of chemical genetics (compared to limited/biased experimental evolution efforts) in mapping CS and XR interactions, and supports that the 103 further drug pair relationships (n=116 total) from our training set warrant reclassification (**Extended Data Fig. 2**).

We started with only 25 CS interactions in the training set (from the 4 published studies). Here we could infer and validate 21 further CS interactions. All of them were monochromatic and the majority (n=15) also bidirectional (**Fig. 3c**). This illustrates the power of chemical genetics to identify new CS interactions, and especially the rare bidirectional ones, which are the most promising for cycling/combination therapies^8–15^. In contrast to CS drug pairs, about a third of the tested XR pairs (n=11/31), including ones previously known, were non-monochromatic (**Fig. 3d)**, i.e. some evolved lineages were sensitive, instead of resistant, to the second antibiotic. In seven XR cases we failed to detect the expected bidirectionality, and in 4 further cases, we failed to detect the interaction overall (**Fig. 3d)**. Overall, this highlights again that experimental evolution experiments are prone to false negative calls (even with large number of lineages being evolved), and uncovers an unexpected tendency for XR interactions to be non-monochromatic.

### Antibiotic classes with extensive XR or CS

In contrast to most other studies looking into CS and XR, where one antibiotic per class is tested, here we could assess more systematically antibiotic class behaviors, as several antibiotic classes were represented by multiple members in the chemical genetics data^31^. As expected, antibiotics belonging to the same class had exclusively XR interactions between them, as they largely share mode of action and mechanisms of resistance. In contrast, as previously shown^39^, antibiotics of different chemical classes, exhibited both XR and CS interactions (**Fig. 4a**), the former often driven by promiscuous resistance mechanisms (e.g. efflux pumps), and the latter by mutations that lead to modifications of the outer membrane composition (**Extended Data Fig. 3**). We next asked whether antibiotic classes behaved coherently, i.e. whether members of two classes interacted predominantly in the same way. Although this was true for antibiotic classes with members that share cellular target(s) and/or transport mechanisms to enter or exit the cell (e.g. tetracyclines, macrolides), this was less of a case for classes with distinct targets (beta-lactams) or distinct transport mechanisms (quinolones of different generations) (**Fig. 4c**). Interestingly, protein synthesis inhibitor classes did not only act coherently, but were also dominated by XR interactions between them (**Fig. 4b**) with the exception of aminoglycosides, which have been reported to show extensive CS interactions with drugs of different classes^8,17,19^.

**Fig 4.**
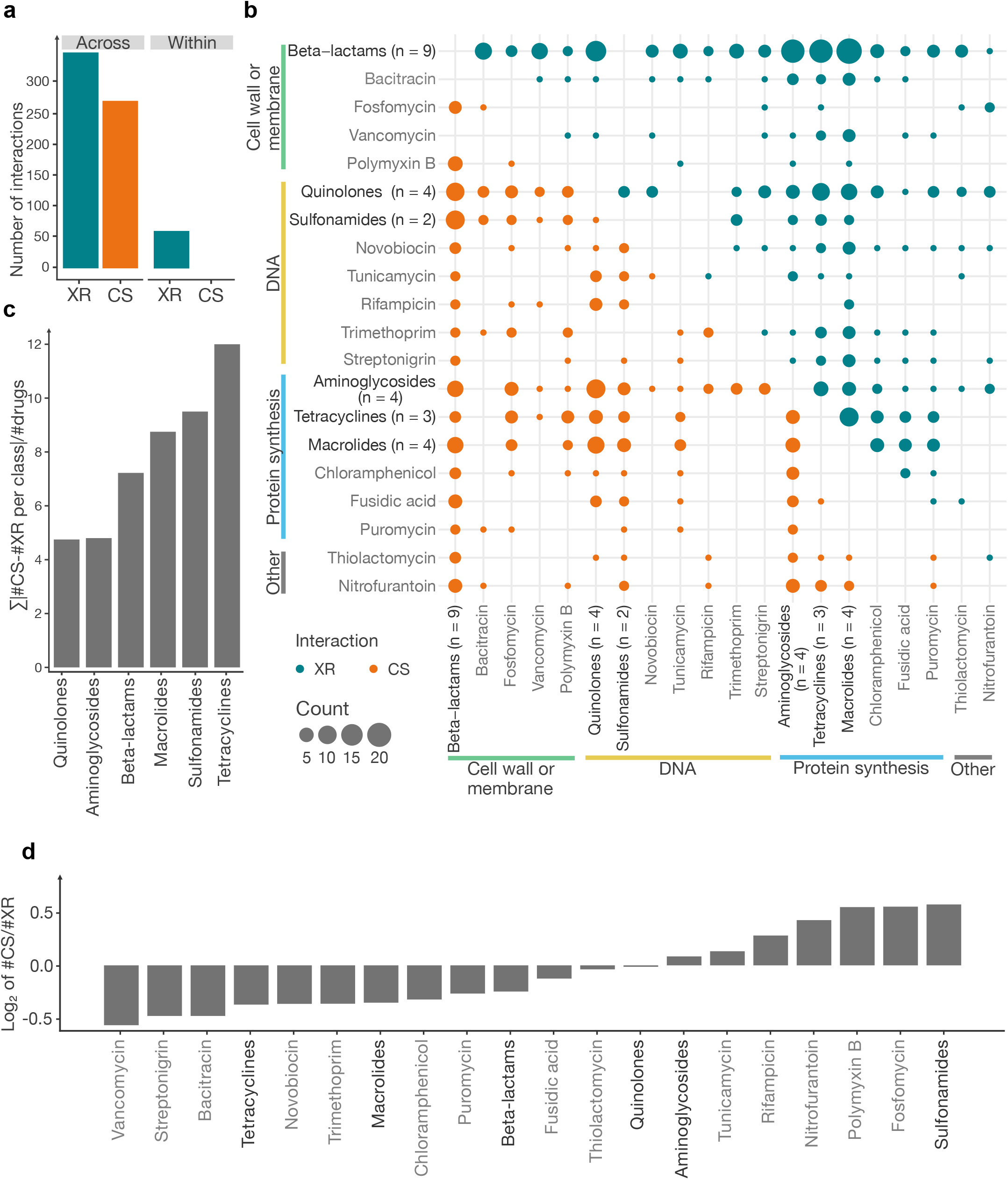
CS and XR interactions between and within antibiotic classes. **a**, Interactions between members of same antibiotic class (within class) are exclusively XR. The within class group includes classes with more than one member: beta-lactams, aminoglycosides, quinolones, macrolides, tetracyclines and sulfonamides. **b,** Overview of all inferred and known drug interactions in *E. coli* at the class level. When a class has only one representative then antibiotic is named and shown in grey. Within class interactions are not displayed in the plot, but are all exclusively XR. Antibiotics are grouped according to their modes of action. Dot size represents the count of interactions between classes (or single antibiotics). **c,** Coherency of interactions of each class with all other classes, calculated by the sum of the absolute differences between XR and CS interactions with each other class, normalized by the number of drugs in the class. The higher the number, the more coherently the class is interacting. **d,** Interaction preference of each class (single- or multi-membered), calculated as the ratio between the number of CS and XR interactions with all other antibiotics from other classes. Antibiotic classes with ratio>1 are considered as predominantly CS (n=8), whereas those with ratio<1 as predominantly XR (n=12).

Besides aminoglycosides, the only other class reported to be enriched in CS interactions are polymyxins^8,17^. In addition to these two classes and nitrofurantoin, which has been reported before^17^, we identified sulfonamides and a number of single drugs (fosfomycin, rifampicin, tunicamycin) with extensive CS interactions (**Fig. 4b, d**). Sulfonamides were largely collateral sensitive to macrolides and beta-lactams, driven by LPS- and nucleotide biosynthesis-related mechanisms (**Fig 4b**, **Extended Data Fig. 3a**). In contrast, protein synthesis inhibitors (apart from aminoglycosides) were enriched in XR interactions, largely because of shared efflux resistance mechanisms (AcrAB-TolC) between them (**Fig 4b, d & Extended Data Fig. 3b**).

### Chemical genetics capture CS and XR mechanisms occurring during evolution

Causal mechanisms behind XR and CS interactions are hard to identify from evolution experiments, as passenger mutations occur in parallel to causal one(s) and indirect mutations can also affect the expression/activity of causal resistance elements. The situation is worse for CS interactions, as very few are known to begin with^7,17,19,26^. Chemical genetics makes it easier to disentangle causality, as all genes contributing to resistance or sensitivity to a certain drug are identified. To prove this point, we first investigated how known CS interactions were represented in chemical genetics. For example, the decrease in proton motive force (PMF) across the inner membrane decreases aminoglycoside uptake and makes cells more resistant to aminoglycosides, but also collateral sensitive to other drugs whose efflux is driven by PMF-dependent pumps, like AcrAB-TolC^17,19^. Mutations in *trkH*, encoding a proton-potassium symporter, were previously shown to cause this phenotype, in particular for the CS interaction between the aminoglycoside tobramycin and nalidixic acid or tetracycline^17,39^. Indeed, the *trkH* mutant, as well as mutants in subunits of the respiratory complexes^17,39^, exhibited extreme discordant s-scores for these known CS drug pairs in chemical genetics (**Extended Data Fig. 4a**). Using the same approach, we tested whether we could unravel the unknown mechanism underpinning the recently described CS interaction between cefoxitin-novobiocin^26^. Genes involved in adding polarity to the lipopolysaccharide (LPS) core, *waaG, waaP and waaQ*, were strongly discordant for this drug pair, leading to cefoxitin resistance and novobiocin sensitivity (**Extended Data Fig. 4b**). The outer membrane (OM) penetration of novobiocin, a large lipophilic antibiotic is known to be affected by LPS modifications^40,41^. At the same time these mutations lower the levels of the OM porins, OmpC and OmpF^42^, allowing cefoxitin and other cephalosporins to enter the cell^43^.

CS and XR interactions can be non-monochromatic, as multiple resistance mechanisms exist for a given drug. Since chemical genetics systematically explore the mutational space, we assumed that they should capture better the dynamics of such interactions. To assess this, we focused on XR drug pairs which exhibited some level of inconsistency in our experimental evolution (n=11/31; **Fig. 3d**). Antibiotic pairs with non-monochromatic XR interactions exhibited significantly stronger discordance scores in chemical genetics than drug pairs with monochromatic XR (**Extended Data Fig. 4c**). Hence chemical genetics can capture monochromaticity of XR interactions, and identify those antibiotic pairs that can also evolve CS relationships (**Extended Data Fig. 4d-g**). We then investigated in more detail the most non-monochromatic pair, that of tetracycline and azithromycin, which showed XR, CS and neutral interactions in 4, 6 and 2 lineages, respectively (**Fig. 3d**). For each of our 12 tetracycline-evolved lineages, we measured changes in susceptibility to both antibiotics at each of the 10 passages (**Fig. 5a**, **Methods**). Almost all lineages exhibited increased neutrality with time, except for three lineages (1, 4 and 12), which evolved the lowest resistance to tetracycline, and remained CS to azithromycin (**Fig. 5a)**. First, this partially explains the low rates of CS and XR discovery in previous studies (**Fig. 2a**), since evolution experiments typically use final populations with high resistance to test for XR/CS. Second, it implies that with time cells evolve more specific resistance mechanisms, e.g. target-compared to intracellular concentration-related ones.

**Fig 5.**
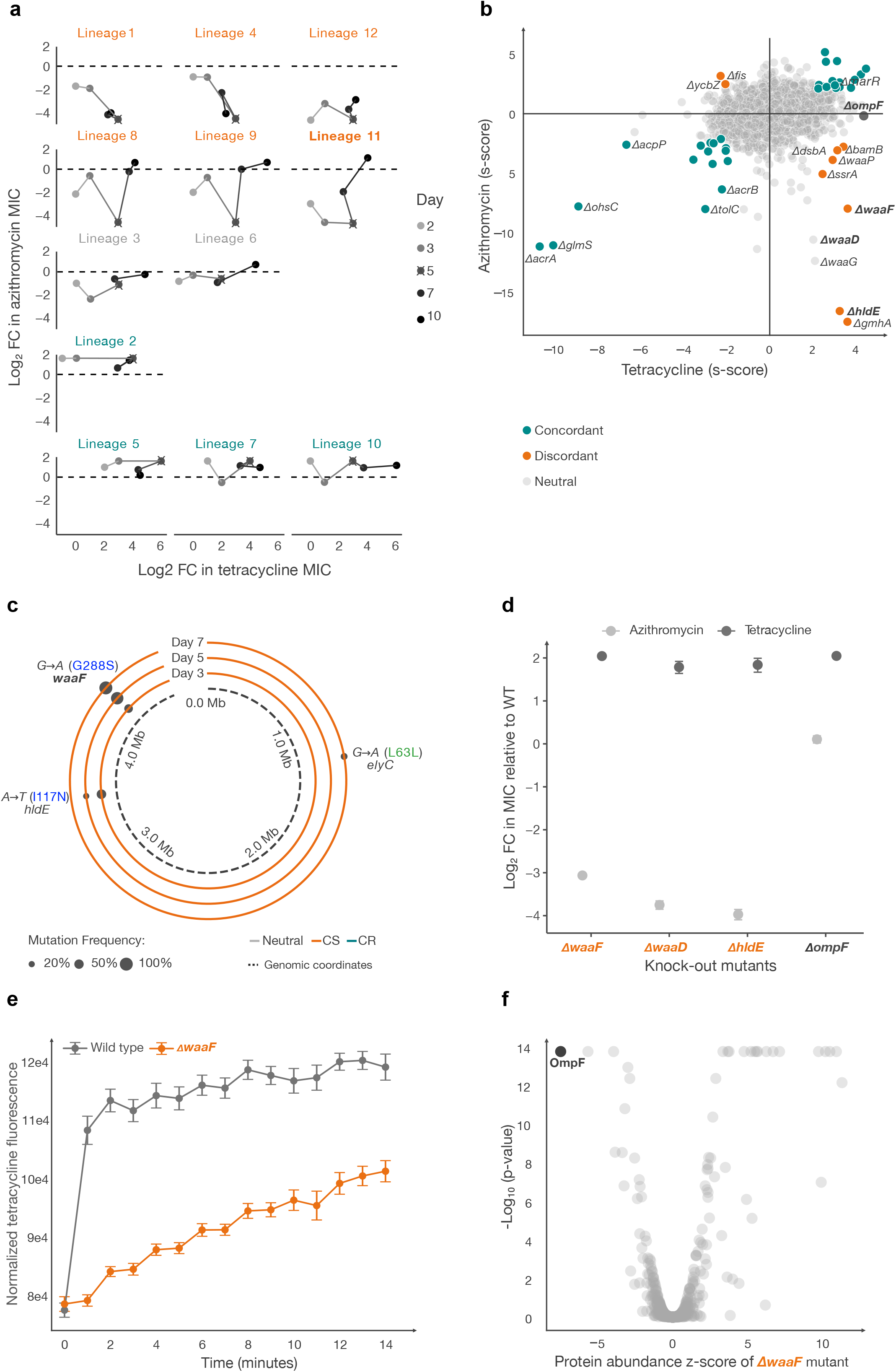
Chemical genetics recapitulates dynamics and explains mechanisms of non-monochromatic interactions. **a**, Changes in azithromycin susceptibility during evolution of 12 lineages in tetracycline. Evolution was performed by passaging every 24 h for 10 days (total 100 generations, **Methods**). Resistance levels of 12 lineages to both antibiotics are shown for days 2, 3, 5, 7 and 10. Lineages are grouped according to whether they exhibited CS, neutrality and XR at day 5 (same as **Fig. 3e**). **b,** Scatter plot of chemical genetic profiles of the *E. coli* deletion library in tetracycline and azithromycin^31^. Mutants with concordant (XR-related) and discordant (CS-related) profiles are highlighted. Dots in grey represent mutants that do not have s-scores within our chosen 3% extreme cutoff for both drugs. **c,** Mutations of lineage 11 during evolution. Genome sequencing of lineage population reveals a succession of two point mutations in genes that both lead to CS - first in *hldE*, which is then replaced by mutations in *waaF*, a slightly less detrimental gene for azithromycin resistance according to chemical genetics data (**b)**. For the other 11 lineages see **Extended Data Fig. 5. d**, Fold changes in tetracycline and azithromycin IC90s of wildtype and knockout mutants confirm that both *hldE* and *waaF* contribute to resistance to tetracycline and sensitivity to azithromycin, while *ompF* deletion leads only to resistance to tetracycline. **e,** Tetracycline uptake is reduced in a *waaF* deletion mutant. Tetracycline fluorescence was measured in cell pellets, and signal was normalized by cellular abundance (OD_600nm_). The mean and standard error are shown (n = 3-6 biological replicates). **f,** OmpF, a major tetracycline importer, is the most downregulated protein in the *waaF* deletion mutant^42^.

To better understand the mechanisms underlying the changes of the tetracycline-azithromycin relationship over time, we sequenced all 12 lineage populations from days 3, 5, and 7 (**Extended Data Fig. 5**). Lineages with neutral interactions carried either point mutations in tetracycline target genes (e.g. lineage 3 with *rpsJ* V57L– coding for the S10 ribosomal protein^44^), or a combination of CS and XR strains in the population (e.g. linage 7 with mutations in *hldE* and *marR*) (**Fig. 5a**, **Extended Data Fig. 5**). Mutations in *marR*, a gene encoding for a repressor of the main transcriptional regulator of efflux pumps in *E. coli* and known modulator of antibiotic resistance^45,46^, were behind all XR interactions observed in different lineages (2, 5, 7 and 10 – **Fig. 5a**, **Extended Data Fig. 5**). This was in agreement with Δ*marR* increased resistance to both drugs in chemical genetics data (**Fig. 5b**). In contrast, all lineages with stable and strong CS interactions had promoter or deletion mutations in *waaD* (**Extended Data Fig. 5**), one of most sensitive mutants to azithromycin and resistant to tetracycline in chemical genetics data^31,47^ (**Fig. 5b**). Lineages that were initially CS but became neutral (8, 9 and 11), carried initially strong CS mutations on *waaD* or *hldE*, both involved in synthesis of the ADP-heptose precursor of core LPS, which were then replaced by strains with mutations in genes with milder CS or XR phenotypes, like *waaF* and *marR* (**Fig. 5b-c**, **Extended Data Fig. 5**). We confirmed the slightly milder CS (lower azithromycin sensitivity) for Δ*waaF*, a gene encoding a protein that adds the second heptose sugar to the LPS inner core, compared to Δ*hldeE or* Δ*waaD* (**Fig. 5d**). We postulated that the increased tetracycline resistance of all these LPS core mutants is due to reduced uptake compared to the wildtype, and confirmed this by measuring intracellular tetracycline fluorescence in Δ*waaF* cells (**Fig. 5e**; **Source Table of Fig. 5e**). This lower intracellular tetracycline is likely due to low OmpF levels in in Δ*waaF* cells (**Fig. 5f**)^42^, as OmpF is the major tetracycline importer^43,48,49^. This is in agreement with chemical genetics data, where Δ*ompF* is tetracycline resistant, but not azithromycin-sensitive (**Fig. 5b, d**). Hence, loss-of-function mutations in *waaF* (or in other LPS core genes, such as *hldE*, *waaD, waaP*) lead to less OmpF in the OM, and tetracycline resistance. At the same time, cells become more sensitive to azithromycin (and macrolides), because the OM becomes less polar and thereby more permeable to hydrophobic antibiotics^50^.

Overall, we confirmed that chemical genetics data can pinpoint CS and XR mechanisms that emerge and get selected during experimental evolution, thereby helping us to even rationalize the dynamics of non-monochromatic antibiotic interactions.

### Combining newly identified CS antibiotic pairs reduces evolution of resistance

The combination, sequential use or cycling of CS drug pairs has been shown to reduce the rate of resistance evolution^8–15^ and re-sensitize resistant strains^16^ in laboratory settings, and for a *Pseudomonas aeruginosa* infection in clinics^23^. Considering the therapeutic potential of CS antibiotic combinations, we tested the degree to which our newly identified CS pairs could reduce resistance evolution in combination, when compared to single drugs (**Fig. 6a**). We selected 4 CS, 2 neutral, and 1 XR pairs involving 9 commonly used antibiotics. For seven parallel *E. coli* lineages, we measured the MIC alone and in combination (using 1:1 ratio compared to drug MICs). We evolved 7 *E. coli* lineages to single drugs or combinations for 7 days, and measured the MIC of the evolved population (**Fig. 6a**, **Methods**). For each antibiotic combination, we calculated 2401 Evolvability Indexes (7^4^ combinations), that is the degree by which resistance to any of the single drugs increases (log_2_ Evolvability Index > 0) or decreases (log_2_ Evolvability Index < 0) in the drug combination (**Methods**)^21^. As expected, lineages evolved in the presence of the ceftazidime-ciprofloxacin XR combination reached higher resistance to each drug, compared to lineages evolved with single antibiotic treatments (**Fig. 6b**; **Source Table of Fig. 6**). In contrast, most lineages treated with CS or neutral combinations evolved lower resistance than those treated with single antibiotics (**Fig. 6b**). The strongest reduction in resistance evolution occurred for combinations of bidirectional CS pairs (**Fig. 3c**, **6b).** For example, 6 out of 7 lineages evolved full resistance towards mecillinam alone (256-fold increase in MIC), while combining mecillinam with nitrofurantoin or levofloxacin led to almost no mecillinam resistance (average fold-change in MIC < 2). For the cefoxitin-levofloxacin pair, resistance evolved in combination was lower just for cefoxitin but not levofloxacin **(Fig. 6b**, **Extended Data Fig. 6**), despite the pair showing bidirectional CS during experimental evolution (**Fig. 3c**). Altogether, we demonstrate that reciprocally CS antibiotic pairs hold a great potential for diminishing resistance evolution when used in combination.

**Fig 6.**
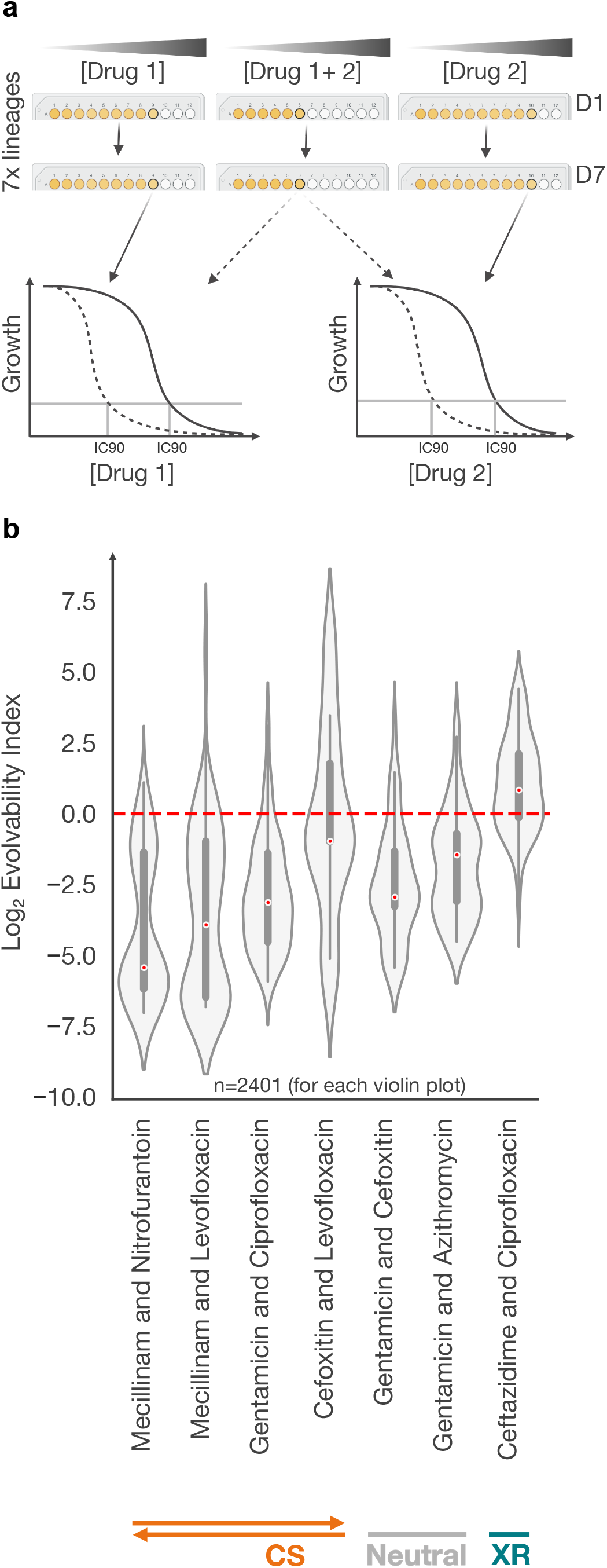
Combinations of reciprocal CS antibiotic pairs reduce resistance evolution. **a**, Experimental design: after evolution of resistance against single antibiotics or their combination (7 lineages for each, passaging every 24h for 7 days, 70 generations in total), the IC90 with both antibiotics was tested for the evolved mutants. In each passage mutants growing (colored as yellow) at highest concentration (denoted by thick circle) were transferred (**Methods**). **b,** Measured IC90 values were used to calculate Evolvability Index (Formula 2; Methods). Red line represents the cutoff (Evolvability Index = 0), below which the antibiotic pair is considered to reduce resistance evolution compared to single antibiotics. Red dots on the violin plots represent the median. Non-XR antibiotic combinations led to lower collective resistance, and in the case of reciprocal CS to lower Evolvability indexes and lower resistance to each of the antibiotics combined (**Extended Data Fig. 6**).

## Discussion

A better understanding of how resistance to one antibiotic limits treatment with others (cross-resistance - XR) or opens new opportunities (collateral sensitivity - CS) is imperative in the context of the ongoing AMR crisis. In the last decade, such drug interactions have been assessed in several pathogens^8,12,16–18,21–25,51^. However, the main detection method, experimental evolution, has obvious limitations. First, it has low sensitivity, which leads to different studies reporting different interactions for the same drug pairs even in the same species (**Fig. 2a**). This is because during experimental evolution experiments, often only a limited number of lineages and resistance mechanisms are probed. What further augments the problem is that resistance mechanisms largely depend on the amount and time of selective pressure applied, as we show here for the tetracycline-azithromycin pair (**Fig. 5a**), and that each study uses different selection pressure, metrics and number of lineages to assess interactions. Although within species comparisons are possible when metric and selection pressure are standardized^52^, cross-species comparisons become prohibitive with high false negative rates. Second, experimental evolution is laborious and limits the number of drug-pairs that can be tested. As a result, monochromaticity of interactions (especially for drug classes) has been impossible to assess properly in the past. Last, it is very hard to identify the underlying mechanism for CS or XR interactions by sequencing the resistance lineages from experimental evolution, and without additional tailored experiments.

By assessing the impact of thousands of individual mutations at once on resistance or sensitivity to different drugs, chemical genetics can bypass most of these limitations. As we show here, chemical genetics offer a way to systematically and quantitatively assess all chromosomal resistance mechanisms (independent of selective pressure), and can dramatically increase the throughput of bacterial species and drugs tested. In addition, it gives insights into how monochromatic, reciprocal or conserved such interactions are, as well as a basis to dissect the driving mechanisms. As proof-of-principle we focused on published chemical genetics data from *E. coli*^31^, because of the large number of antibiotics screened at different concentrations and the extensive benchmarking. In the future, similar analyses can be expanded to other available datasets in the same or other species^34,47,53–56^. Such datasets will inevitably increase with time, as genome-wide mutant libraries are becoming available in tens of species and even more strains^57,58^, whether those are arrayed or pooled^29,59^, and constructed by targeted deletions^60–62^, transposon insertions^59,63^, or CRISPRi knockdowns^53,64^. Including such libraries will allow to probe the role of essential genes and/or gene overexpression when mapping antibiotic resistance and XR/CS relationships.

In this study we devised a new approach and metric to map CS and XR in *E. coli*, using available chemical genetics data for 40 antibiotics. Thereby, we increased the number of known interactions by several-fold, and resolved more than a hundred cases of prior conflicts and/or misclassifications reported in literature. Beyond this, we obtained unique insights into within-class interactions, unraveling that all antibiotic classes are dominated by XR interactions between their members. Although this is largely expected, some classes have members with non-overlapping targets and/or resistance mechanisms. Specifically for beta-lactams, their use in combination has been reported to constrain resistance evolution, during fast switching regimens^65^ or for specific pairs and resistance mechanisms^66^. Moreover, we identified many new bidirectional CS interactions, and used a handful to show that evolution of antibiotic resistance against combinations of such antibiotics is harder. Last, we mechanistically rationalized CS interactions and explained why some drug interactions can be non-monochromatic. In the case of tetracycline-azithromycin, the mechanisms that played a role in experimental evolution were a small subset of the possible mechanisms revealed by chemical genetics. This is likely due to probing only 12 lineages, but also likely due to the fitness costs of some of these resistance mechanisms. Interestingly, the interaction changed non-monotonically over time, and longer/stronger selection on one drug (tetracycline) led to more neutral interactions to the second one (azithromycin). This means that long-term, bacterial populations may opt for target mutations or low/neutralized fitness cost resistance mechanisms, neutralizing also CS/XR interactions. Hence fast switching or combinatorial treatments may be more efficient than sequential antibiotic treatments for CS drug pairs.

In the future, the increased ability to map XR and CS interactions between drugs opens the path for expanding such endeavors to non-antibiotics with antibacterial or adjuvant activity^67–69^, and to probing interactions in different environments, such as in bile, different pH^70^, urine media, biofilms^71^ or gut microbiome communities, as fitness costs are known to change with environment^72^. Moreover, the systematic nature of chemical genetics limits false negatives and metric biases, and can allow for comprehensive comparisons across species and strains using corresponding genome-wide mutant libraries. Cross-species studies have been conducted previously to map drug synergies and antagonisms^35,73^. Knowing how drugs interact at multiple levels - resistance evolution, efficacy (growth inhibition or killing), long-term clearance effects^74^, and host cytotoxicity will open the path for designing more effective and long-lasting combinations for clinics.

## Materials and Methods

### Data sources and preprocessing

The *E. coli* chemical genetics data were obtained from a previous study^31^, where the fitness of 3979 non-essential single-gene knockout mutants and essential gene hypomorphs was evaluated in 324 different conditions (114 unique stresses and drugs tested in different concentrations). Fitness effects were quantified as s-scores, i.e. a modified t-statistic on the deviation of the colony size of one mutant in one condition from the median colony size of the mutant across all conditions^38,75^. We reprocessed the data to exclude: a) strains from the hypomorphic mutant collection and mutants that had 10 or more missing values for the conditions - reaching a final number of 3904 mutants; and b) environmental stresses (e.g. different temperatures, pH, heavy metals, amino acids, dyes and alternative carbon sources), non-antibiotic drugs, and drug combinations. Antibiotics with a narrow range of s-scores (no extreme s-scores below -6.9 or above 3.9) were also excluded from analysis (n=7). This left us with 40 antibiotics that were further used in this study (**Supplementary Table 1**). For those antibiotics tested in multiple concentrations, the highest one was selected.

Previously reported XR and CS interactions were collected from four studies. Viktoria Lazar *et al.*^17,18^ measured XR and CS in *E. coli* BW25113 using 12 antibiotics where interactions were defined based on at least a 10% difference in the growth of more than 50% evolved lineages compared to control lineages. Tugce Oz *et al*.^19^ and Lejla Imamovic and Morten O. A. Sommer^8^ compared MICs of evolved populations against the wildtype to define XR and CS in *E. coli* MG1655 using 22 and 23 antibiotics respectively. We kept the original definitions and assessments of XR and CS used in the respective study. When integrating these datasets, interactions of overlapping antibiotic pairs were annotated as “XR & Neutral”, “CS & Neutral”, “XR & CS”, and “XR & CS & Neutral” if conflicting interactions were observed in different studies. Interactions with “XR & CS” and “XR & CS & Neutral” annotations were removed (n=6) and “XR & Neutral” and “CS & Neutral” were reannotated as “XR” and “CS”, respectively, because evolution experiments are prone to false negatives. Directionality was reduced (keeping drug 1 - drug 2 but removing reciprocal) by removing one pair (if XR/CS was bidirectional) or by removing the “neutral pair” (if the interaction was unidirectional). After the preprocessing steps, only conditions for which chemical genetics data was available were selected as training set (n=24), amounting to 111 neutral, 70 XR, and 25 CS drug pair relationships (**Supplementary Table 3)**.

### Assessment of correlation metrics

Since the first attempts of combining chemical genetics profiles and XR/CS interactions found associations between chemical genetics profile similarity and XR/CS^17,18^, we assessed several correlation methods from SciPy^76^ to compute various correlation coefficients between two drugs (Drug 1 and Drug 2; **Extended Data Fig 1a**). The correlation functions were applied to drug pairs with known interactions for which chemical genetics data is available. For each drug pair in this dataset, the correlation coefficient was computed for the four methods (Pearson, Spearman, Kendall Tau, and Weighted Tau). Receiver Operating Characteristic (ROC) curves were plotted to evaluate the performance of the computed correlation coefficients in distinguishing between interaction types (XR (n=70) vs non-XR (n=136), and CS (n=25) vs non-CS (n=181)). The correlation coefficients served as the predictor values, and the interaction types (either XR or CS) were the true labels. The area under the ROC curve (AUC) was computed for each correlation method (**Extended Data Fig. 1a**).

### Feature generation and interpretation of decision trees

For each condition in the chemical genetic data, 3% extreme positive and negative s-scores were chosen after assessment of different cutoffs (**Extended Data Fig. 1d**). Six features were generated by antibiotics pairwise calculation: sum of positive concordant s-scores, sum of negative concordant s-scores, sum of discordant s-scores, count of positive concordant s-scores, count of negative concordant s-scores, and count of discordant s-scores. Using these features, machine learning algorithms (based on decision trees^77^) were used and models were trained to classify XR (n=70) vs non-XR (n=136) and CS (n=25) vs non-CS (n=181).

To address the class imbalance, the minority class was oversampled to match the size of the majority class. A search space for hyperparameters was defined for the decision tree classifier, including the function to measure the quality of a split, the maximum depth of the tree, the minimum number of samples required to split an internal node, and the minimum number of samples required to be at a leaf node. A five-fold grid search cross-validation, stratified to maintain the same proportion of the target class as the entire dataset, was used to find the best hyperparameters for the decision tree classifier based on the F1 score. The resulting classifier was trained and again evaluated on the balanced dataset using cross-validation. The best classifier according to F1 score, precision, recall, and ROC AUC was then fitted to the balanced dataset.

The trained decision tree classifier was visualized, showing the decision paths and splits. The tree visualization was limited to a depth of 3 for clarity (**Extended Data Fig. 1c**). We learned from decision tree classifiers that if the count of concordant negative s-scores was higher, the level of discordance was not important to classify interactions. The sum and count of concordant negative s-scores were found to be the most important features, followed by the sum of discordant s-scores. This information was used to generate the OCDM metric, described in detail below. Classifier training, hyperparameter tuning, and visualization were implemented using the scikit-learn package (version 1.1.3)^78^.

### Metric generation and interaction measurement

Among correlation methods, six chemical genetics derived features, and their engineered combinations, we identified the outlier concordance-discordance difference metric (OCDM) as the best metric to statistically significantly separate XR, neutral and CS interactions (**Fig. 2c**). OCDM is defined as the difference between the sum of concordant s-scores and the sum of discordant s-scores if the count of concordant s-scores (N_C_) is below the median count as shown below. Otherwise, OCDM is simply the sum of concordant s-scores.

**Formula 1:**

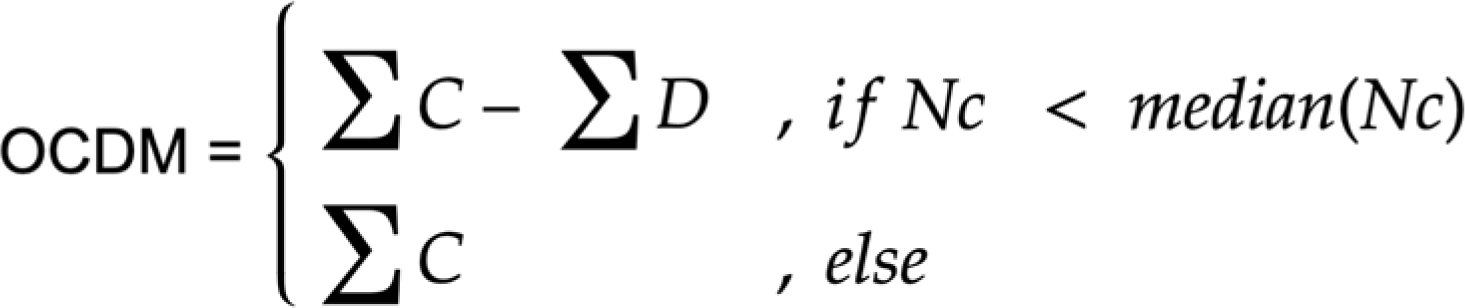

where C represents concordant s-scores and D represents discordant s-scores. To identify optimal threshold determination (cutoffs) of OCDM, False Positive Rate (FPR) and True Positive Rate (TPR) were used to calculate True Factor (TF) = TPR−(1−FPR) = Sensitivity - Specificity, which was computed for each threshold. This threshold represents the best trade-off between sensitivity (TPR) and specificity (1-FPR), which are >105.159057 (to define XR) and <27.224792 (to define CS).

All data analysis was performed in Python (v3.9.17).

### Bacterial strains and growth conditions

For all experiments and unless otherwise specified, *E. coli* (strain BW25113) was grown in LB Lennox broth (tryptone 10 g l−1, yeast extract 5 g l−1, sodium chloride 5 g l−1) at 37°C and fully aerobically (850 rpm), or on agar (2%) plates (same media and temperature).

### MIC determination

*E. coli* BW25113 overnight cultures were diluted to an OD_600nm_ of 0.001 and grown with antibiotics (**Supplementary Table 1**) at eight concentrations on a two-fold dilution gradient, in two technical replicates in microtiter plates (U bottom 96-well plates, Greiner Bio-One 268200) at 37 °C with continuous shaking (850 rpm - orbital microplate shaking). Plates were sealed with breathable membrane (Breathe-Easy; Z380059-1PAK) and OD_600nm_ was measured every 30 mins for 24 hours. The liquid handler Biomek FX (Beckman Coulter) was used to prepare plates. All MIC tests were performed in a total volume of 100 µL per well.

Controls included: no-cell + no-drug controls to assess contamination, no-drug controls to assess maximal growth, no-cell controls to assess artifacts (OD_600nm_ change) of the drugs alone or of their interaction with medium components. The area under the growth curve was calculated using simps function from SciPy^76^ and divided by the no-drug control. 90% inhibitory concentration (IC90 which we define as MIC) was calculated using the drc package in R^79^.

### Experimental evolution and XR/CS measurements

*E. coli* wildtype overnight cultures were diluted 1:1000 and exposed to 23 antibiotics in eight concentrations from 0.5 x IC90 to 64 x IC90, in 12 lineages using the same volumes and plates as for MIC determination. Every 24 hours, the lineages growing in the highest concentration (OD_600nm_ > 0.3) were back-diluted to OD_600nm_ of 0.01, and the volume needed to reach a final dilution of 1:1000 (3-10 µL) was transferred to the next plate with the same concentration gradients. Once the evolution experiment was completed (5 passages for total of 5 days; ∼50 generations in total), the lineages were tested for antibiotic susceptibility for 59 of the 634 predicted interactions (9.3%; 23 novel XR, 8 known XR, 21 novel CS, 2 known CS, 4 novel neutral, and 1 known neutral interaction; **Fig. 3c-e** & **Source Table of Fig. 3**). MICs were measured as in “MIC determination” (12 lineages/populations x 118 drug pairs (59 unique drug pairs) x 2 technical replicates = 2832 MIC values; **Source Table of Fig. 3c-e**). Changes in IC90 were compared to the ancestor strain. Interactions were defined as XR or CS if log_2_ fold-change > +1 or -1, respectively. For the azithromycin-tetracycline pair, we performed five more passages (total of 10 passages; ∼100 generations), and tracked changes both in tetracycline resistance and azithromycin susceptibilities.

### Whole-genome sequencing and analysis

A clone from the wildtype and from populations of 12 lineages from day 3, 5, and 7 were sequenced to determine mutations responsible for given phenotype. Genomic DNA was extracted using Macherey Nagel DNA extraction kit and sequenced using single-end Illumina NextSeq 2000 (P1; length of 122 bp). Mutations were identified by mapping sequences to the reference genome from the NCBI database (*E. coli* BW25113 strain K-12 chromosome; GCF_000750555.1)^80^ using Breseq ^81^ with the following parameters: -p -l 80 -j 8 -b 5 -m 30. Mutations present in the wildtype clone compared to the NCBI reference genome were eliminated to only identify mutations that are associated with resistance/sensitivity.

### P1 transduction

Single colonies of *E. coli* wildtype (BW25113) and corresponding Keio mutants^60^ were used for P1 transduction. P1 lysate preparation and transduction was performed as previously described^82^. We confirmed the transduction success via colony PCR.

### Tetracycline fluorescence assay

Wildtype and knockout mutants of *waaF, waaD,* and *hldE,* and *E. coli* was grown in 5 mL LB with continuous shaking at 37°C until they reached an OD_600nm_ of 0.5. 1 mL aliquots of each culture were centrifuged at 3,500 rpm for 10 minutes and supernatants were discarded. Pellets were further washed three times with 0.5 mL of 137 mM PBS, and resuspended in 50 µL of 137 mM PBS and transferred in black-walled, clear- and flat-bottom 96-well plates (Greiner Bio-One 655096), containing three concentrations of two-fold serially diluted tetracycline (highest final concentration 16 µg/mL, final volume 100 µL/well). OD_600 nm_ and fluorescence (excitation λ 405 nm and emission λ 535 nm) were measured with an Infinite M1000 PRO plate reader (Tecan), for 15 minutes with readings taken every minute. Experiments were conducted for three to six biological replicates.

### Experimental evolution against antibiotic combinations

IC90 for individual antibiotics (n=8) and drug combinations at 1:1 MIC ratio (n=7) were measured as in “MIC determination”. The evolution experiment was carried out in the same way as described in “Experimental evolution and XR/CS measurements” with the following changes: the initial wildtype culture was exposed to 8 single and 7 combinations of antibiotics in 11 concentrations from 0.125 x IC90 to 128 x IC90, for 7 lineages. At the end of the experiment (7 passages for the total of 7 days; ∼70 generations), IC90 of drug 1 and 2 were measured in drug 1, drug 2 and drug 1 + drug 2 resistant lineages as described in “MIC measurements”. To compare evolution of resistance to single drugs vs drug combinations, Evolvability Indexes were calculated for each possible pair (2401 values per antibiotic combination) as shown below.

**Formula 2:**

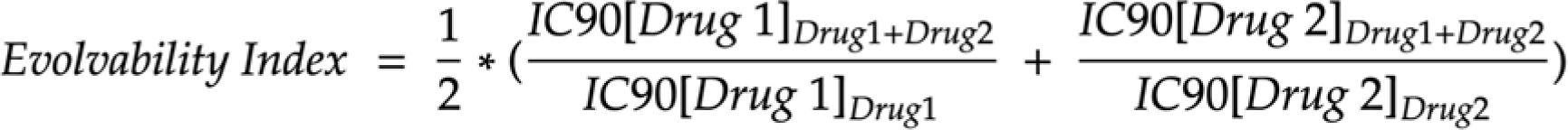

where IC90[Drug 1]_Drug1 + Drug2_ corresponds to IC90 of drug 1 for lineage evolved against drug 1+2 combination.

**Extended Data Fig. 1. Performance of different metrics & models in capturing XR and CS antibiotic interactions from chemical genetics data. a.** Receiver operating characteristic (ROC) curves for classification of XR (positive class) vs non-XR (negative class), and CS (positive class) vs non-CS (negative class), using simple linear and non-linear correlation metrics. AUC is the area under the curve. **b.** Performance of the decision tree model on balanced classes shows that both XR and CS interactions can be well classified. **c.** decision tree with classes CS (class 1) versus the rest (class 0), where maximum depth of 3 is shown for visualization, illustrates hierarchy of decisions to discriminate classes. Each node in the tree represents a decision point based on the value of a particular feature, and branches represent the outcome of the decision. The root node divides the data based on the concordant_negative_w feature, which is a sum of s-scores (as weights) of hits on negative concordant site of a scatterplot. The tree branches out to discordant_w feature, which is a sum of s-scores (as weights) of hits on discordant site of a scatterplot, while discordant_w_m is a sum of products of s-scores (as weights) of hits on discordant site of a scatterplot. **d.** P-values from a paired Mann-Whitney U-test (two-sided) are depicted across quantile cutoffs for extreme s-scores to differentiate XR/CS/neutral interactions based on OCDM values. Q3 and Q97 perform the best.

**Extended Data Fig. 2. Chemical genetics metric captures well prior information and reclassifies a subset of prior interactions. a-b.** Comparison of previously reported XR (**a**) and CS (**b**) interactions with our inferences based on our chemical genetics metric (OCDM) show an agreement of 67-68% for CS (n=17) and XR (n=47) - 11 such interactions were validated experimentally during our benchmarking (**Fig. 3c-d**). The rest is wrongly inferred as neutral or the opposite interaction, including four interactions (3 XR & 1 CS) that we experimentally validated as false positives (**Fig. 3c-e**). **c.** In contrast to CS or XS, there is less agreement for neutral interactions with previous studies. This is consistent with the high false negative rates when comparing prior studies between them (**Fig. 2a**). The majority of previously reported neutral interactions (76.6%, n=85) are inferred as CS/XR by chemical genetics. All 8 we included in the benchmarking set were confirmed as false negatives (**Fig. 3c-e**). **d.** New XR, Neutral and CS pairs inferred by chemical genetics and the OCDM cutoff are 2.8- and 6.4-fold more that currently known XR and CS antibiotic interactions in *E. coli*, after reclassifying interactions (n = 116) we infer differently than previously reported. This plot includes interactions that are known and for which chemical genetics data is not available.

**Extended Data Fig. 3. Chemical genetics can uncover the biological processes that drive interactions between antibiotic classes. a.** Clustered heatmap of discordant mutants that are part of CS interactions between sulfonamides and macrolides (blue) or beta-lactams (green). Genes in bold are involved in LPS or nucleotide biosynthesis. **b.** Clustered heatmap of concordant mutants that are part of XR interactions between tetracyclines (violet), macrolides (blue) and other protein synthesis inhibitors. Genes in bold regulate or are part of the major efflux pump in *E. coli* (AcrAB-TolC).

**Extended Data Fig. 4. Chemical genetics can infer mechanisms and monochromaticity of XR and CS drug interactions. a.** Scatter plot of chemical genetic profiles of the *E. coli* deletion library in tobramycin and nalidixic acid^31^. Mutants with concordant (XR-related) and discordant (CS-related) profiles are highlighted. Dots in grey represent mutants that do not have s-scores within the 3% extreme values for both drugs. The underlined knockout mutants are known causal genes of this CS interaction^17,19^. **b.** Chemical genetic profiles for novobiocin and cefoxitin, presented as in **a**. Underlined knockout mutants indicate that the changes in polarity of the lipopolysaccharide (LPS) core can drive resistance to cefoxitin, while providing sensitivity to the large and non-polar novobiocin. **c.** Non-monochromatic XR interactions (n=11) have higher absolute discordance scores than their monochromatic counterparts (n=20) (Mann-Whitney U-test) - monochromaticity was defined in the validation experiment. This means that chemical genetics can infer monochromaticity of XR interactions. **d**, Highest discordance score of -133.8481 based on the 11 non-monochromatic XR interactions from **c** was used to separate all XR interactions into monochromatic (n=230) or non-monochromatic (n=174). **e-g.** Scatter plots of chemical genetic profiles of the *E. coli* deletion library^31^ for examples of other pairs of drugs with both high concordance and discordance (in addition to azithromycin and tetracycline shown in **Fig. 5b**). As the azithromycin-tetracycline pair, those are expected to be non-monochromatic. Data are depicted as in **a**-**b**.

**Extended Data Fig. 5. Genome sequencing of lineage populations evolved in tetracycline.** Results of remaining 11 lineages from days 3, 5, and 7. Results shown as in **Fig. 5c**, and lineages grouped in XR, CS and neutral according to classification in **Fig. 5a**.

**Extended Data Fig. 6. Antibiotics combinations constrain resistance evolution to both, one or none of the compounds.** The log2 of MIC (IC90) of evolved population in both drugs compared evolved population to drug itself is used to identify whether and how well combining drugs reduces resistance to each drug compared to single-drug treatments. Reciprocal CS drug pairs do this efficiently. Red dashed line shows the no-effect, when combining drugs is not changing resistance evolution to single drug treatments.

## Acknowledgements

We thank Marco Galardini, Alexandra Koumoutsi, and the EMBL Gene Core for help with sequencing experiments; Alexandra Koumoutsi, Luo Yan Yong and Stefan Bassler for experimental advice; the Typas lab and especially Karin Mitosch for fruitful discussions. This work was supported by the ERC consolidator grant (uCARE) to A.T.

## Author contributions

A.T. and N.S. conceived and designed the study. A.T., C.G., E.C., P.B. and J.M. supervised the project. All scripts were written by N.S., with advice on data pre-processing from F.H., on machine learning from A.O., and MIC determination from V.V. All experiments were carried out by N.S. with advice from E.C and C.G. Figures were designed and plotted by N.S. with inputs from E.C, A.O, C.G, and A.T. N.S. and A.T. wrote the manuscript with input from all authors. All authors approved the final version.

## Competing Interest declaration

Authors have no competing interests to declare.

## Author information

Correspondence and requests for materials should be addressed to &

## Notes

### Competing Interest Statement

The authors have declared no competing interest.

## References

1. Murray, C. J. L. et al. Global burden of bacterial antimicrobial resistance in 2019: a systematic analysis. The Lancet 399, 629–655 (2022).

2. Theuretzbacher, U. et al. Analysis of the clinical antibacterial and antituberculosis pipeline. Lancet Infect Dis 19, e40–e50 (2019).

3. Butler, M. S., Henderson, I. R., Capon, R. J. & Blaskovich, M. A. T. Antibiotics in the clinical pipeline as of December 2022. J Antibiot (Tokyo) 76, 431–473 (2023).

4. Szybalski, W. & Bryson, V. Genetic Studies on Microbial Cross-Resistance to Toxic Agents I. J Bacteriol 64, 489–499 (1952).

5. Beutner, E. H., Doyle, W. M. & Evander, L. C. Collateral Susceptibility of Isoniazid-Resistant Tubercle Bacilli To Nitrofurans. American Review of Respiratory Disease (1963).

6. Baym, M., Stone, L. K. & Kishony, R. Multidrug evolutionary strategies to reverse antibiotic resistance. Science 351, (2016).

7. Roemhild, R., Linkevicius, M. & Andersson, D. I. Molecular mechanisms of collateral sensitivity to the antibiotic nitrofurantoin. PLoS Biol 18, e3000612 (2020).

8. Imamovic, L. & Sommer, M. O. A. Use of Collateral Sensitivity Networks to Design Drug Cycling Protocols That Avoid Resistance Development. Science Translational Medicine 5, 204ra132–204ra132 (2013).

9. Kim, S., Lieberman, T. D. & Kishony, R. Alternating antibiotic treatments constrain evolutionary paths to multidrug resistance. Proc. Natl. Acad. Sci. U.S.A. 111, 14494– 14499 (2014).

10. Barbosa, C., Beardmore, R., Schulenburg, H. & Jansen, G. Antibiotic combination efficacy (ACE) networks for a Pseudomonas aeruginosa model. PLOS Biology 16, e2004356 (2018).

11. Roemhild, R. et al. Cellular hysteresis as a principle to maximize the efficacy of antibiotic therapy. Proc Natl Acad Sci U S A 115, 9767–9772 (2018).

12. Hernando-Amado, S., Sanz-García, F. & Martínez, J. L. Rapid and robust evolution of collateral sensitivity in Pseudomonas aeruginosa antibiotic-resistant mutants. Science Advances 6, eaba5493 (2020).

13. Aulin, L. B. S. et al. Design principles of collateral sensitivity-based dosing strategies. Nat Commun 12, 5691 (2021).

14. Jahn, L. J. et al. Compatibility of Evolutionary Responses to Constituent Antibiotics Drive Resistance Evolution to Drug Pairs. Molecular Biology and Evolution 38, 2057–2069 (2021).

15. Hernando-Amado, S., Laborda, P. & Martínez, J. L. Tackling antibiotic resistance by inducing transient and robust collateral sensitivity. Nat Commun 14, 1723 (2023).

16. Barbosa, C., Römhild, R., Rosenstiel, P. & Schulenburg, H. Evolutionary stability of collateral sensitivity to antibiotics in the model pathogen Pseudomonas aeruginosa. Elife 8, (2019).

17. Lázár, V. et al. Bacterial evolution of antibiotic hypersensitivity. Mol. Syst. Biol. 9, 700 (2013).

18. Lázár, V. et al. Genome-wide analysis captures the determinants of the antibiotic cross-resistance interaction network. Nat Commun 5, 4352 (2014).

19. Oz, T. et al. Strength of Selection Pressure Is an Important Parameter Contributing to the Complexity of Antibiotic Resistance Evolution. Mol Biol Evol 31, 2387–2401 (2014).

20. Arcangioli, M.-A., Leroy-Setrin, S., Martel, J.-L. & Chaslus-Dancla, E. Evolution of chloramphenicol resistance, with emergence of cross-resistance to florfenicol, in bovine Salmonella Typhimurium strains implicates definitive phage type (DT) 104. Journal of Medical Microbiology, 49, 103–110 (2000).

21. Rodriguez de Evgrafov, M., et al. Collateral Resistance and Sensitivity Modulate Evolution of High-Level Resistance to Drug Combination Treatment in Staphylococcus aureus. Mol. Biol. Evol. 32, 1175–1185 (2015).

22. Barbosa, C. et al. Alternative Evolutionary Paths to Bacterial Antibiotic Resistance Cause Distinct Collateral Effects. Mol Biol Evol 34, 2229–2244 (2017).

23. Imamovic, L. et al. Drug-Driven Phenotypic Convergence Supports Rational Treatment Strategies of Chronic Infections. Cell 172, 121–134.e14 (2018).

24. Laborda, P., Martínez, J. L. & Hernando-Amado, S. Convergent phenotypic evolution towards fosfomycin collateral sensitivity of Pseudomonas aeruginosa antibiotic-resistant mutants. Microbial Biotechnology 15, 613–629 (2022).

25. Hernando-Amado, S., Laborda, P., Valverde, J. R. & Martínez, J. L. Mutational background influences P. aeruginosa ciprofloxacin resistance evolution but preserves collateral sensitivity robustness. Proceedings of the National Academy of Sciences 119, e2109370119 (2022).

26. Liu, D. Y. et al. Collateral sensitivity profiling in drug-resistant Escherichia coli identifies natural products suppressing cephalosporin resistance. Nat Commun 14, 1976 (2023).

27. Suzuki, S., Horinouchi, T. & Furusawa, C. Prediction of antibiotic resistance by gene expression profiles. Nature Communications 5, 5792 (2014).

28. Horinouchi, T. et al. Prediction of Cross-resistance and Collateral Sensitivity by Gene Expression profiles and Genomic Mutations. Scientific Reports 7, 14009 (2017).

29. Brochado, A. R. & Typas, A. High-throughput approaches to understanding gene function and mapping network architecture in bacteria. Curr Opin Microbiol 16, 199–206 (2013).

30. Cacace, E., Kritikos, G. & Typas, A. Chemical genetics in drug discovery. Current Opinion in Systems Biology 4, 35–42 (2017).

31. Nichols, R. J. et al. Phenotypic Landscape of a Bacterial Cell. Cell 144, 143–156 (2011).

32. Ezraty, B. et al. Fe-S cluster biosynthesis controls uptake of aminoglycosides in a ROS-less death pathway. Science 340, 1583–1587 (2013).

33. Chandrasekaran, S. et al. Chemogenomics and orthology-based design of antibiotic combination therapies. Mol Syst Biol 12, 872 (2016).

34. Shiver, A. L. et al. A Chemical-Genomic Screen of Neglected Antibiotics Reveals Illicit Transport of Kasugamycin and Blasticidin S. PLoS Genet 12, e1006124 (2016).

35. Brochado, A. R. et al. Species-specific activity of antibacterial drug combinations. Nature 559, 259–263 (2018).

36. Silvis, M. R. et al. Morphological and Transcriptional Responses to CRISPRi Knockdown of Essential Genes in Escherichia coli. mBio 12, e0256121 (2021).

37. Kintses, B. et al. Chemical-genetic profiling reveals limited cross-resistance between antimicrobial peptides with different modes of action. Nat Commun 10, 1–13 (2019).

38. Collins, S. R., Schuldiner, M., Krogan, N. J. & Weissman, J. S. A strategy for extracting and analyzing large-scale quantitative epistatic interaction data. Genome Biol 7, R63 (2006).

39. Pál, C., Papp, B. & Lázár, V. Collateral sensitivity of antibiotic-resistant microbes. Trends in Microbiology 23, 401–407 (2015).

40. Møller, A. K. et al. An Escherichia coli MG1655 lipopolysaccharide deep-rough core mutant grows and survives in mouse cecal mucus but fails to colonize the mouse large intestine. Infect Immun 71, 2142–2152 (2003).

41. Nobre, T. M. et al. Modification of Salmonella Lipopolysaccharides Prevents the Outer Membrane Penetration of Novobiocin. Biophys J 109, 2537–2545 (2015).

42. Mateus, A. et al. The functional proteome landscape of Escherichia coli. Nature 588, 473–478 (2020).

43. Mortimer, P. G. & Piddock, L. J. The accumulation of five antibacterial agents in porin-deficient mutants of Escherichia coli. J Antimicrob Chemother 32, 195–213 (1993).

44. Hu, M., Nandi, S., Davies, C. & Nicholas, R. A. High-level chromosomally mediated tetracycline resistance in Neisseria gonorrhoeae results from a point mutation in the rpsJ gene encoding ribosomal protein S10 in combination with the mtrR and penB resistance determinants. Antimicrob Agents Chemother 49, 4327–4334 (2005).

45. Grkovic, S., Brown, M. H. & Skurray, R. A. Regulation of bacterial drug export systems. Microbiol Mol Biol Rev 66, 671–701, table of contents (2002).

46. Beggs, G. A., Brennan, R. G. & Arshad, M. MarR family proteins are important regulators of clinically relevant antibiotic resistance. Protein Sci 29, 647–653 (2020).

47. Price, M. N. et al. Mutant phenotypes for thousands of bacterial genes of unknown function. Nature 557, 503–509 (2018).

48. Cohen, S. P. et al. Cross-resistance to fluoroquinolones in multiple-antibiotic-resistant (Mar) Escherichia coli selected by tetracycline or chloramphenicol: decreased drug accumulation associated with membrane changes in addition to OmpF reduction. Antimicrob Agents Chemother 33, 1318–1325 (1989).

49. Thanassi, D. G., Suh, G. S. & Nikaido, H. Role of outer membrane barrier in efflux-mediated tetracycline resistance of Escherichia coli. J Bacteriol 177, 998–1007 (1995).

50. Nikaido, H. Molecular Basis of Bacterial Outer Membrane Permeability Revisited. Microbiol Mol Biol Rev 67, 593–656 (2003).

51. Yen, P. & Papin, J. A. History of antibiotic adaptation influences microbial evolutionary dynamics during subsequent treatment. PLOS Biology 15, e2001586 (2017).

52. Podnecky, N. L. et al. Conserved collateral antibiotic susceptibility networks in diverse clinical strains of Escherichia coli. Nat Commun 9, 1–11 (2018).

53. Peters, J. M., et al. A Comprehensive, CRISPR-based Functional Analysis of Essential Genes in Bacteria. Cell 165, 1493–1506 (2016).

54. Johnson, E. O. et al. Large-scale chemical-genetics yields new M. tuberculosis inhibitor classes. Nature 571, 72–78 (2019).

55. Liu, H. et al. Functional genetics of human gut commensal Bacteroides thetaiotaomicron reveals metabolic requirements for growth across environments. Cell Rep 34, 108789 (2021).

56. Shiver, A. L. et al. A mutant fitness compendium in Bifidobacteria reveals molecular determinants of colonization and host-microbe interactions. bioRxiv 2023.08.29.555234 (2023) doi:10.1101/2023.08.29.555234.

57. Rosconi, F. et al. A bacterial pan-genome makes gene essentiality strain-dependent and evolvable. Nat Microbiol 7, 1580–1592 (2022).

58. Rousset, F. et al. The impact of genetic diversity on gene essentiality within the Escherichia coli species. Nat Microbiol 6, 301–312 (2021).

59. Voogdt, C. G. P. et al. Randomly barcoded transposon mutant libraries for gut commensals II: Applying libraries for functional genetics. Cell Rep 43, 113519 (2023).

60. Baba, T. et al. Construction of Escherichia coli K-12 in-frame, single-gene knockout mutants: the Keio collection. Mol Syst Biol 2, 2006.0008 (2006).

61. Porwollik, S. et al. Defined Single-Gene and Multi-Gene Deletion Mutant Collections in Salmonella enterica sv Typhimurium. PLoS One 9, e99820 (2014).

62. Koo, B.-M. et al. Construction and Analysis of Two Genome-Scale Deletion Libraries for Bacillus subtilis. Cell Systems 4, 291–305.e7 (2017).

63. Tripathi, S. et al. Randomly barcoded transposon mutant libraries for gut commensals I: Strategies for efficient library construction. Cell Rep 43, 113517 (2023).

64. de Bakker, V., Liu, X., Bravo, A. M. & Veening, J.-W. CRISPRi-seq for genome-wide fitness quantification in bacteria. Nat Protoc 17, 252–281 (2022).

65. Batra, A. et al. High potency of sequential therapy with only β-lactam antibiotics. Elife 10, e68876 (2021).

66. Rosenkilde, C. E. H. et al. Collateral sensitivity constrains resistance evolution of the CTX-M-15 β-lactamase. Nat Commun 10, 618 (2019).

67. Wright, G. D. Antibiotic Adjuvants: Rescuing Antibiotics from Resistance. Trends Microbiol 24, 862–871 (2016).

68. Maier, L. et al. Extensive impact of non-antibiotic drugs on human gut bacteria. Nature 555, 623–628 (2018).

69. Tyers, M. & Wright, G. D. Drug combinations: a strategy to extend the life of antibiotics in the 21st century. Nat Rev Microbiol 17, 141–155 (2019).

70. Allen, R. C., Pfrunder-Cardozo, K. R. & Hall, A. R. Collateral Sensitivity Interactions between Antibiotics Depend on Local Abiotic Conditions. mSystems 6, e01055–21 (2021).

71. Santos-Lopez, A. et al. Evolutionary pathways to antibiotic resistance are dependent upon environmental structure and bacterial lifestyle. Elife 8, e47612 (2019).

72. Björkman, J. et al. Effects of environment on compensatory mutations to ameliorate costs of antibiotic resistance. Science 287, 1479–1482 (2000).

73. Cacace, E. et al. Systematic analysis of drug combinations against Gram-positive bacteria. Nat Microbiol 8, 2196–2212 (2023).

74. Lázár, V., Snitser, O., Barkan, D. & Kishony, R. Antibiotic combinations reduce Staphylococcus aureus clearance. Nature 610, 540–546 (2022).

75. Kritikos, G. et al. A tool named Iris for versatile high-throughput phenotyping in microorganisms. Nat Microbiol 2, 1–10 (2017).

76. Virtanen, P. et al. SciPy 1.0: fundamental algorithms for scientific computing in Python. Nat Methods 17, 261–272 (2020).

77. Breiman, L. Classification and Regression Trees. (Routledge, New York, 2017). doi:10.1201/9781315139470.

78. Pedregosa, F. et al. Scikit-learn: Machine Learning in Python. Journal of Machine Learning Research 12, 2825–2830 (2011).

79. Ritz, C., Baty, F., Streibig, J. C. & Gerhard, D. Dose-Response Analysis Using R. PLOS ONE 10, e0146021 (2015).

80. Sayers, E. W. et al. Database resources of the National Center for Biotechnology Information. Nucleic Acids Res 50, D20–D26 (2021).

81. Deatherage, D. E. & Barrick, J. E. Identification of mutations in laboratory evolved microbes from next-generation sequencing data using breseq. Methods Mol Biol 1151, 165–188 (2014).

82. Thomason, L. C., Costantino, N. & Court, D. L. E. coli Genome Manipulation by P1 Transduction. Current Protocols in Molecular Biology 79, 1.17.1-1.17.8 (2007).

