## Supplementary figures and images for "Systematic mapping of antibiotic cross-resistance and collateral sensitivity with chemical genetics"

### Extended Data Fig. 1

Extended Data Figure 1

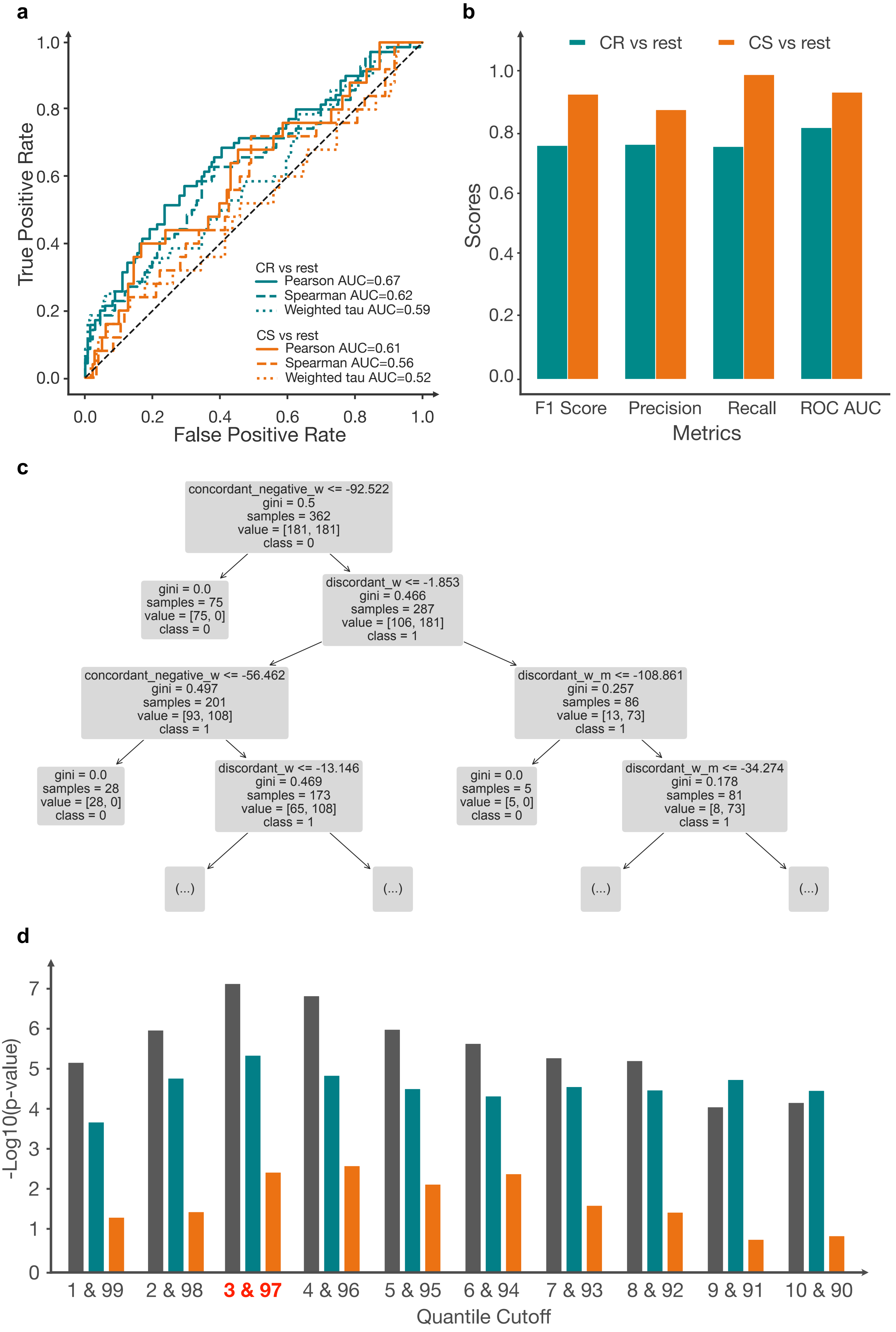

### Extended Data Fig. 2

Extended Data Figure 2

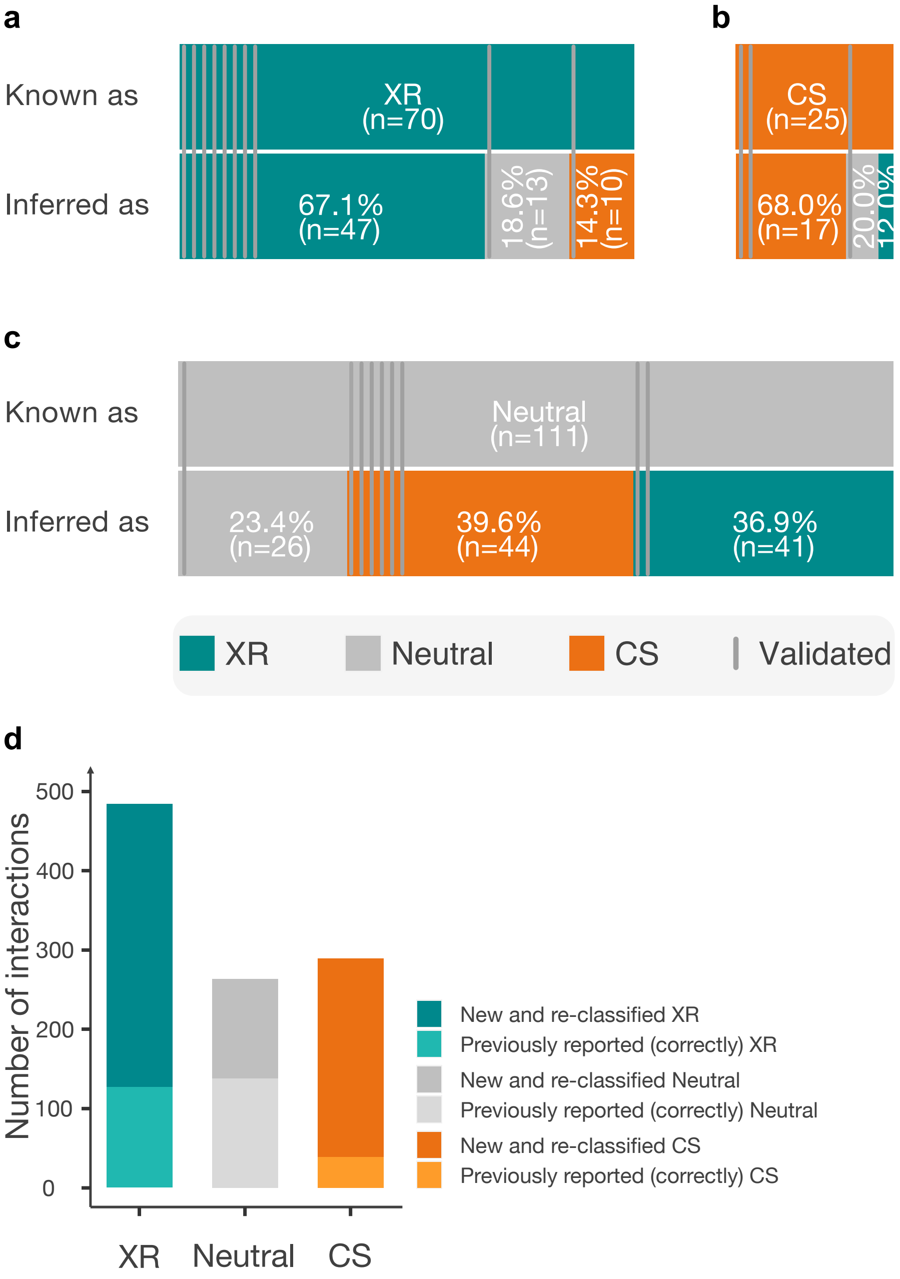

### Extended Data Fig. 3

Extended Data Figure 3

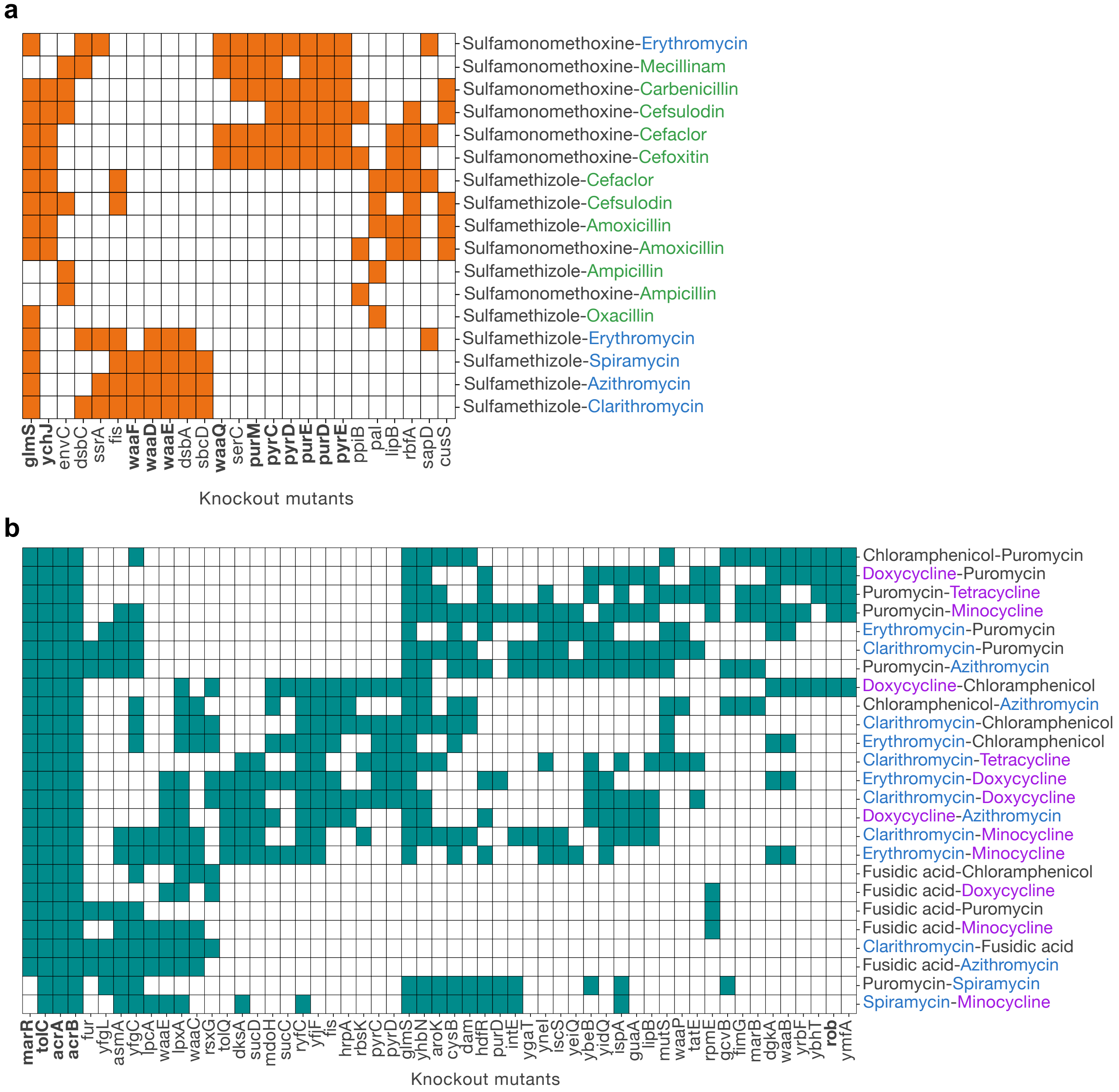

### Extended Data Fig. 4

Extended Data Figure 4

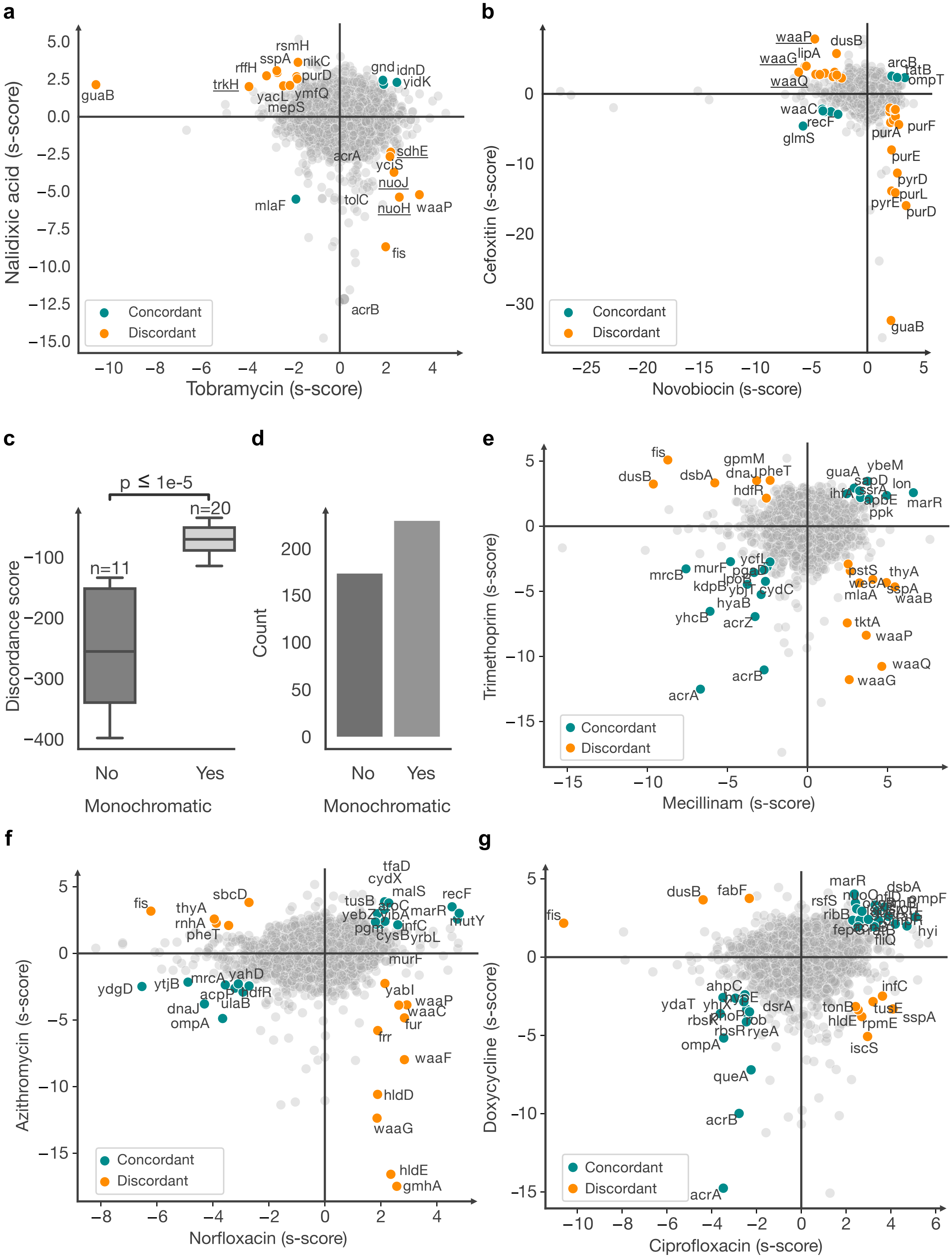

### Extended Data Fig. 5

Extended Data Figure 5

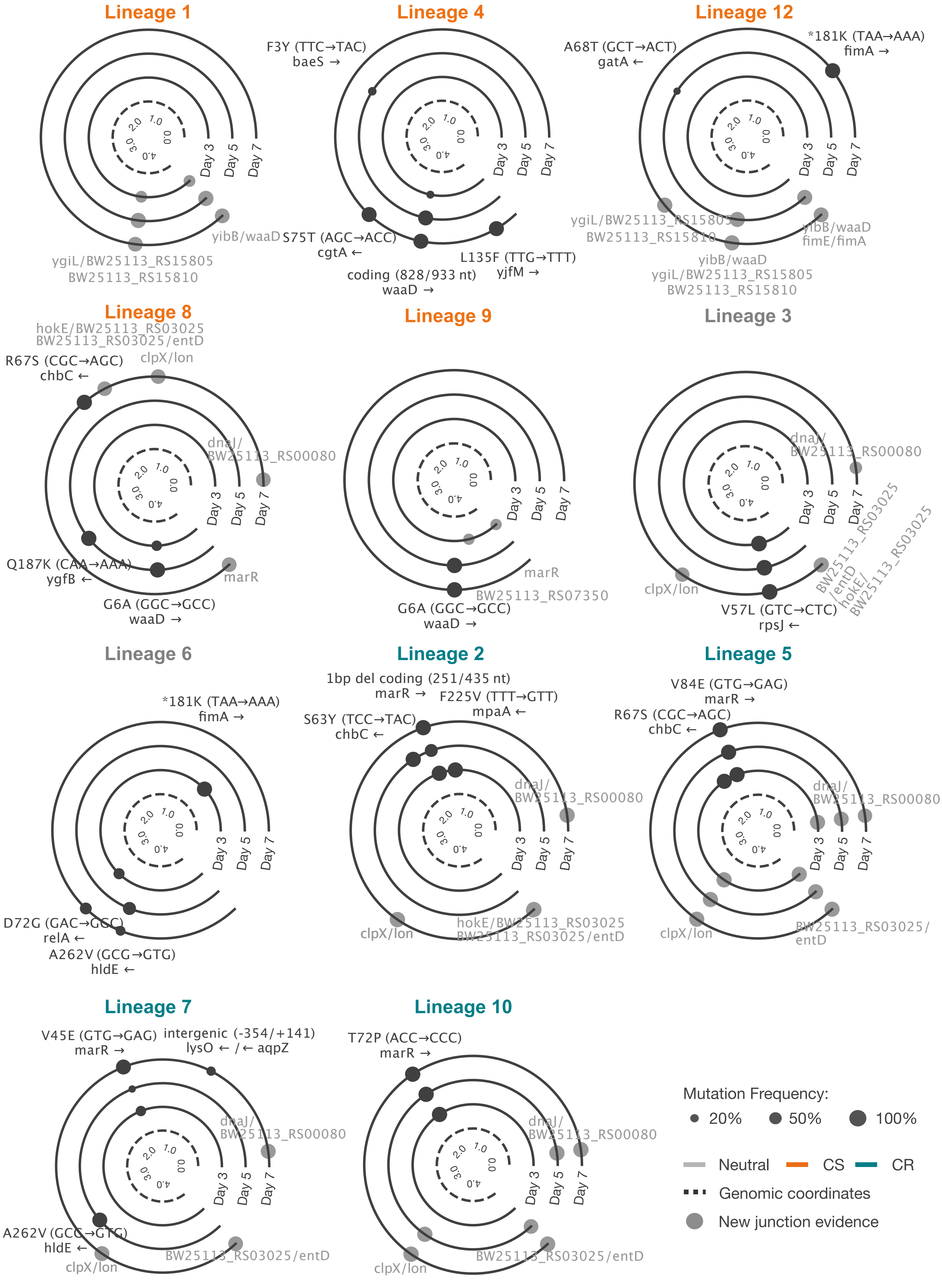

### Extended Data Fig. 6

Extended Data Figure 6

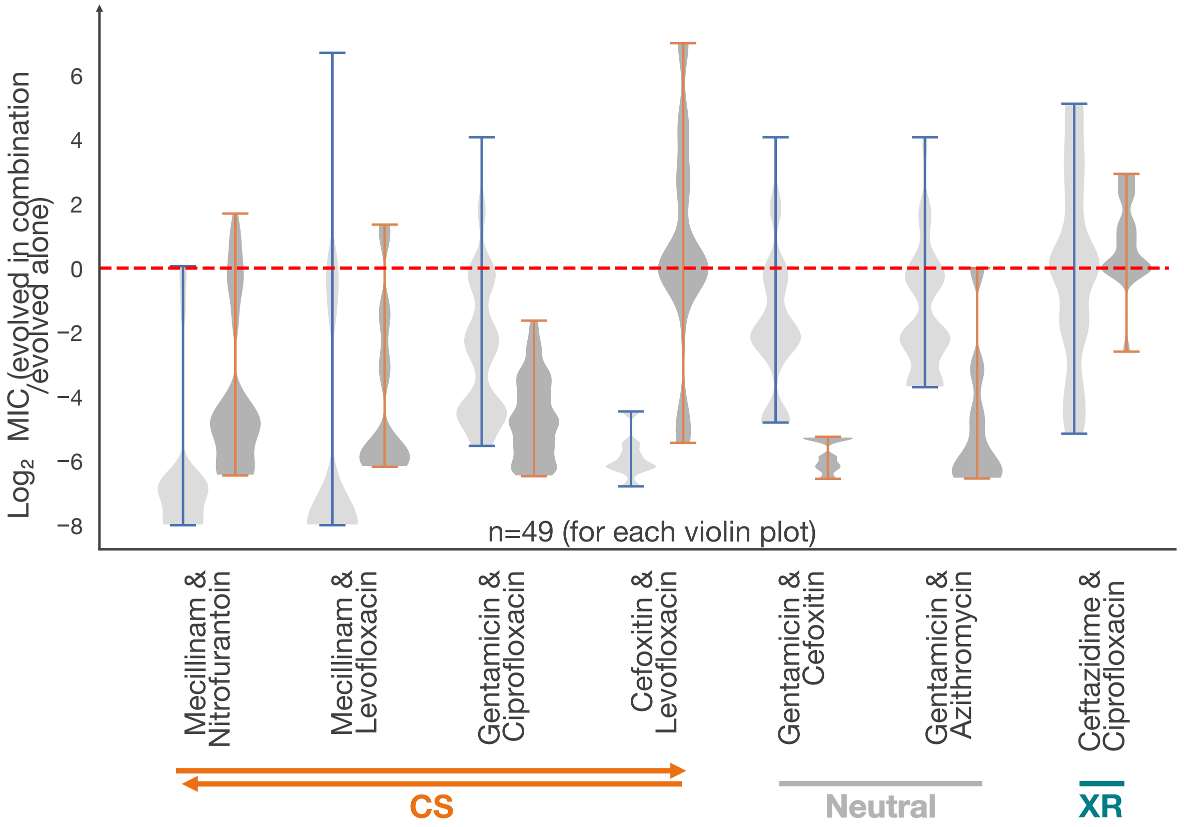
